# A burst of genetic innovation in actin-related proteins (Arps) for testis-specific function in a *Drosophila* lineage

**DOI:** 10.1101/665299

**Authors:** Courtney M. Schroeder, John Valenzuela, Glen M. Hocky, Harmit S. Malik

**Author notes:** Correspondence to be addressed to: Harmit S. Malik, 1100 Fairview Ave N A2-025, Seattle, WA 98109.

## Abstract

Many cytoskeletal proteins form the core of fundamental biological processes and are evolutionarily ancient. For example, the superfamily of actin-related proteins (Arps) specialized early in eukaryotic evolution for diverse cellular roles in the cytoplasm and the nucleus. Despite its strict conservation across eukaryotes, we find that the Arp superfamily has undergone dramatic lineage-specific diversification in *Drosophila*. Our phylogenomic analyses reveal four independent Arp gene duplications that originated in the common ancestor of the *obscura* group of *Drosophila* species and have been mostly preserved in this lineage. All four Arp paralogs have evolved under positive selection and are predominantly expressed in the male germline. We focus our analyses on the divergent *Arp2D* paralog, which arose via a retroduplication event from *Arp2*, a component of the 7-membered Arp2/3 complex that polymerizes branched actin networks. Computational modeling analyses suggest that Arp2D should be able to replace Arp2 in the Arp2/3 complex and bind daughter actin monomers, suggesting that Arp2D may augment Arp2’s functions in the male germline. We find that Arp2D is expressed during and following meiosis in the male germline, where it localizes to distinct locations such as actin cones–specialized cytoskeletal structures that separate bundled spermatids into individual mature sperm. We hypothesize that this unprecedented burst of genetic innovation in cytoskeletal proteins may have been driven by the evolution of sperm heteromorphism in the *obscura* group of *Drosophila*.

## Introduction

Actin is one of the most evolutionarily conserved proteins in eukaryotes; the actin fold even pre-dates eukaryotes and is found among polymerizing proteins in bacteria (van den Ent, et al. 2001) and archaea (Izore, et al. 2016). The canonical structure of actin is necessary for numerous architectural and signaling roles in eukaryotes, including cell-shape maintenance, cell motility, vesicle transport and cytokinesis (Kabsch, et al. 1990; Dominguez and Holmes 2011). The utilitarian actin fold is also conserved in actin-related proteins (Arps) (Kabsch, et al. 1990; Frankel and Mooseker 1996; Dominguez and Holmes 2011). Eight conserved subfamilies of Arps specialized early in eukaryotic evolution for diverse cellular roles in the cytoplasm and the nucleus. These functions include facilitating the polymerization of actin (Arp2 and Arp3) (Mullins, et al. 1998), promoting the motility of the microtubule-based motor dynein (Arp1 and Arp10), (Muhua, et al. 1994; Lee, et al. 2001) and participating in chromatin remodeling (Arps 4,5,6,8) (Harata, et al. 2000; Blessing, et al. 2004; Klages-Mundt, et al. 2018). Of the Arp superfamily members, only Arp1 can form actin-like filaments (Schafer, et al. 1994), whereas most Arps function in complex with other proteins (Machesky, et al. 1994; Klages-Mundt, et al. 2018).

Previous phylogenetic analyses have highlighted the evolution and distinguishing features of different Arp families (Goodson and Hawse 2002; Muller, et al. 2005). Using Arp sequences collected predominantly from five model organisms (*Arabidopsis thaliana, Saccharomyces cerevisiae, Caenorhabditis elegans, Drosophila melanogaster*, and *Homo sapiens*), these studies defined the Arp subfamilies with phylogenetic precision (Goodson and Hawse 2002; Muller, et al. 2005). While Arp subfamilies diverged from each other early in eukaryotic evolution, the Arp sequences within each subfamily are highly similar, indicating the strong selective pressure for conservation of both sequence and function for each Arp (Goodson and Hawse 2002; Muller, et al. 2005). Despite the compelling conservation of cytoplasmic and nuclear Arps across eukaryotic species, studies have also uncovered cases of lineage-specific gains and losses. For example, Arp1 and Arp10 were lost in plants most likely due to the concomitant loss of the motor dynein (Hammesfahr and Kollmar 2012). Similarly, the Arp7 and Arp9 subfamilies are evolutionary inventions specific to fungi (Cairns, et al. 1998; Peterson, et al. 1998; Goodson and Hawse 2002).

Establishing the framework of well-defined Arp lineages has also aided in the identification of lineage-specific “orphan” Arps that do not fall into a defined Arp subfamily (Goodson and Hawse 2002). For example, mammals have approximately 7 testis-specific Arps with no known ortholog outside mammals. Whereas the highly conserved canonical Arps (1-10) are ubiquitously expressed, these ‘orphan’ Arps are primarily expressed in mammalian male germ cells, where some are implicated in spermatogenesis and fertility (Heid, et al. 2002; Tanaka, et al. 2003; Hara, et al. 2008; Boeda, et al. 2011; Fu, et al. 2012). ‘Orphan’ Arp lineages have also been previously described in *Drosophila*. For example, all sequenced species of Drosophilids encode *Arp53D*, which has no orthologous Arp gene outside insects (Goodson and Hawse 2002). Intriguingly, like the mammalian ‘orphan’ Arps, *Arp53D* is also testis-specific in *D. melanogaster* and *D. pseudoobscura* (Fyrberg, et al. 1994; Celniker, et al. 2009), although its functional role remains unknown. Beyond the phylogenomic and expression studies, testis-specific ‘orphan’ Arps have received little scrutiny, presumably due to absence of orthologs in most phyla. Studies of such ‘orphan’ Arps could reveal how a strikingly conserved superfamily can diversify in sequence to innovate cellular roles.

Here, we took a phylogenomic approach to uncover additional innovation in the Arp superfamily in *Drosophila*. Using a comprehensive survey of the Arp superfamily in 12 sequenced and well-annotated *Drosophila* species (Drosophila 12 Genomes 2007), we sought to determine if there were additional Arp paralogs beyond *Arp53D*. Unexpectedly, we discovered 4 lineage-specific Arp paralogs, all of them occurring in the common ancestor of the *obscura* clade of *Drosophila*. Despite the convergence of Arp innovation in the same *Drosophila* lineage, we find that the Arp paralogs arose independently, via duplications of distinct parental Arps or actins. Most of these *obscura*-specific Arps have been retained over the 14 million-year-old lineage, where they evolve under positive selection. Similar to *Arp53D*, we find that all *obscura*-specific Arps are expressed in the testis. Cytological analyses reveal that these Arps may specialize for gametic actin structures, such as motile actin cones that act during sperm individualization. Our results reveal that a burst of genetic innovation in the conserved Arp superfamily allowed for specialized roles in spermatogenesis in a specific lineage of *Drosophila*.

## Results

### A burst of lineage-specific duplications in the *obscura* group of Drosophila

We performed a phylogenomic survey of the *Drosophila* Arp superfamily in the 12 sequenced and well-annotated *Drosophila* genomes (Fig. 1A) (Drosophila 12 Genomes 2007). In addition to several members of the cytoplasmic (canonical) actin gene family, *D. melanogaster* encodes eight Arps (*Arp1-6,8,10*) and one ‘orphan’ *Arp53D* (Fyrberg, et al. 1994; Goodson and Hawse 2002). We used the protein sequences of all nine of these *D. melanogaster* Arps in tBLASTn searches of the 12 sequenced and annotated *Drosophila* species (Drosophila 12 Genomes 2007). All hits with highly significant E-values were collected, aligned, and subjected to phylogenetic and shared syntenic analysis (Fig. 1B). Our analyses revealed that most sequences were orthologs of the previously identified Arp subfamilies. However, we found four additional Arp homologs that were present in *D. pseudoobscura* and *D. persimilis*, two closely-related species that diverged ∼ 1 million years ago (Babcock and Anderson 1996). These Arp paralogs (named Dup1, Dup2, Dup3 and Arp2D) are phylogenetically distinct from the highly conserved canonical Arp subfamilies (Fig. 1B).

**Fig. 1:**
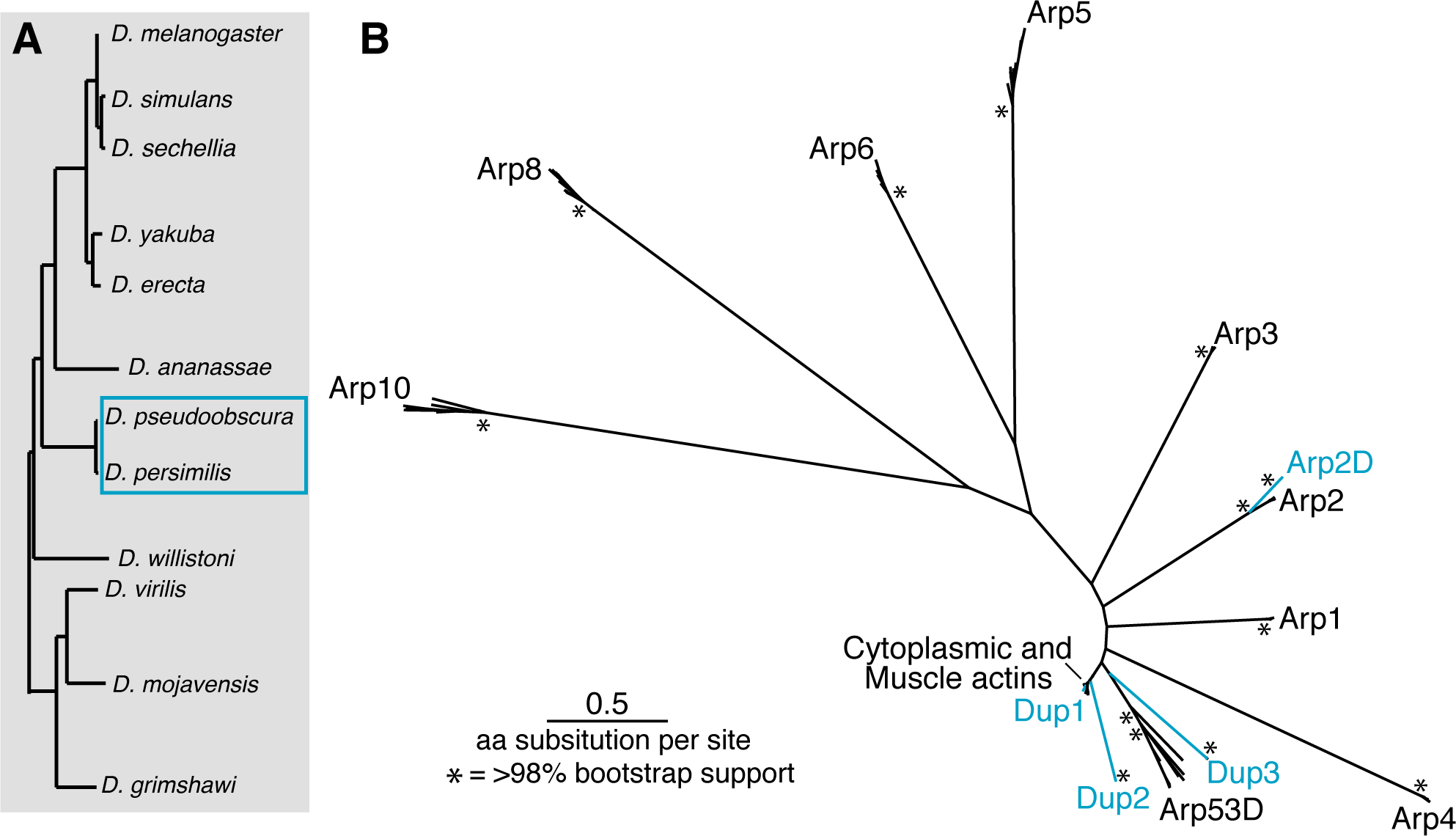
The Arp superfamily exhibits lineage-specific duplications. **A)** *D. melanogaster* Arps were used in a tBLASTn search in the sequenced and annotated genomes of the twelve species displayed. Gene duplications in (B) were found in the two species outlined by a blue box. **B)** All unique hits with E-value of ∼0 were translated and aligned using MAFFT (Katoh and Standley 2013). Gaps where less than 80% of sequences aligned were removed. A PhyML tree (Guindon, et al. 2010) with 100X resampling was generated (LG substitution rate). The asterisks represent nodes with greater than 98% bootstrap support. The highly conserved “canonical” Arps are labeled in black, whereas the blue-labeled branches represent Arp paralogs only found in *D. pseudoobscura* and *D. persimilis*.

We performed an analysis of shared synteny to confirm that these newly identified Arp paralogs are specific to *D. pseudoobscura* and *D. persimilis*. We used the genomic locus of each of the 4 novel *D. pseudoobscura* Arp paralogs to identify shared syntenic locations on each of the other 10 sequenced, annotated *Drosophila* genomes. In each case, we were able to confirm that the Arp genes were indeed missing in the shared syntenic location of species other than *D. pseudoobscura* and *D. persimilis* (Supplementary Fig. S1, S2). Therefore, it appears that all four Arp paralogs arose in one lineage of *Drosophila*, which includes *D. pseudoobscura* and *D. persimilis*.

To further pinpoint the evolutionary age and origin of the four Arp paralogs, we extended our analyses to other members of the *obscura* group, which consists of a number of species that have a common ancestor originating ∼14 mya (Barrio and Ayala 1997; Russo 2013). Using the alignment of the syntenic loci in *D. melanogaster* and one or more *obscura* group species’ draft genomes (Mia Levine, personal communication), we designed primers to locations of high conservation in genes or intergenic regions neighboring each of the four novel Arp paralogs. Using these primers, we performed PCR-based targeted-sequencing of the Arp paralogs in 10 additional species of the *obscura* group whose genome sequences have not been sequenced (Fig. 2, Supplementary Data S1). Phylogenetic analyses based on nucleotide alignments of the genes and pseudogenes from each group recapitulated expected relationships between the *obscura* species (Supplementary Fig. S3), thus confirming our identification of true orthologs of the ‘new’ Arp genes found in *D. pseudoobscura*.

**Fig. 2:**
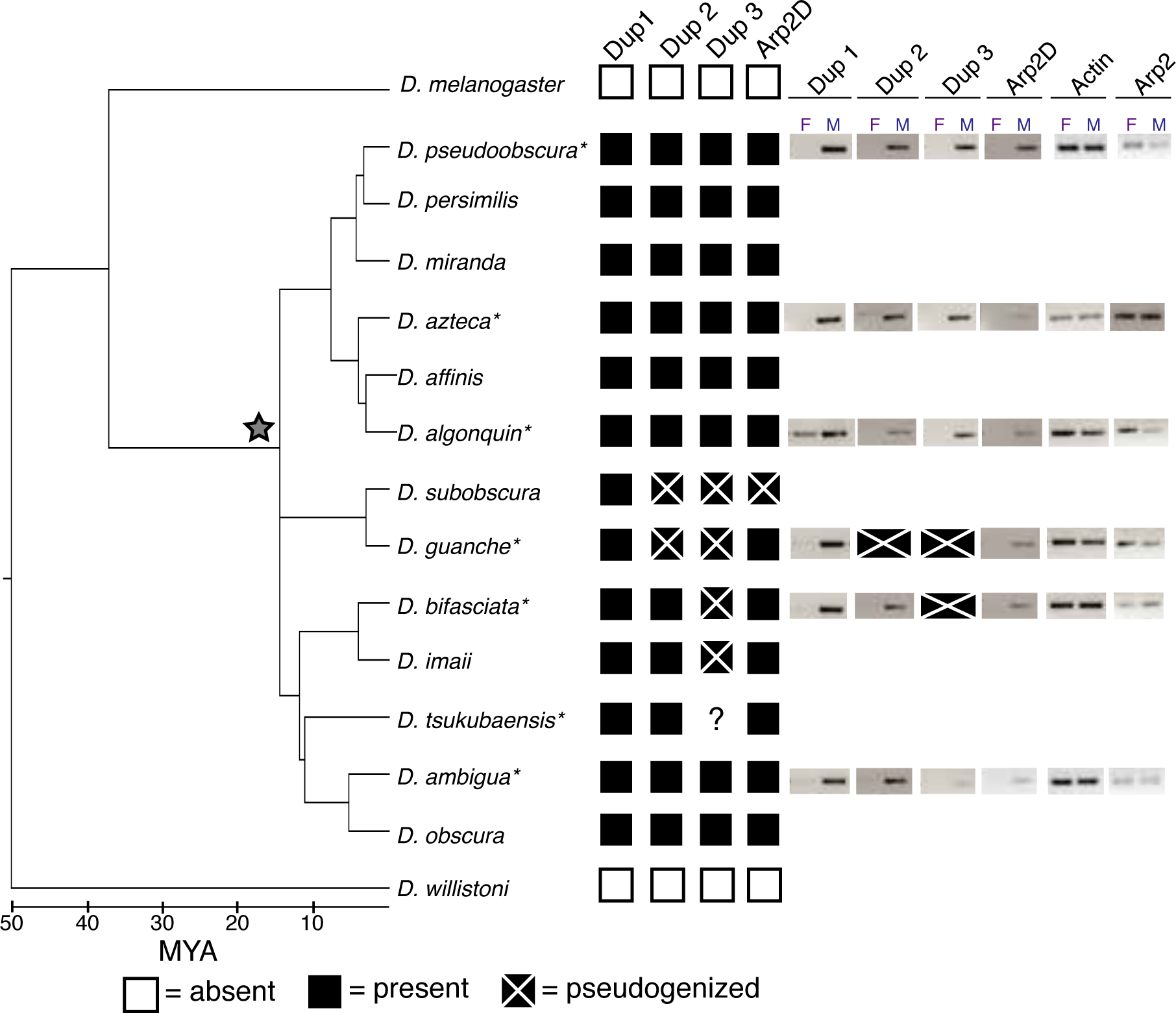
Four Arp paralogs originated in the common ancestor of the *obscura* group and maintained male-specific expression. Shared syntenic loci of the four Arp duplicates were PCR-amplified from 12 species in the *obscura* clade. The presence and absence of the duplicates are indicated with black and white boxes, respectively. Pseudogenized genes are indicated with boxes that have an “X,” and a star indicates the origin of the Arp duplicates. PCR amplification of the *Dup3* locus from *D. tsukubaensis* was unsuccessful so its status is unknown. RT-PCR was conducted with males and females of representative species in the *obscura* clade and DNA gels are displayed. Expression of canonical *actin (Act5C)* and *Arp2* are shown in the gray box. *Dup2* and/or *Dup3* were excluded from the analysis for *D. guanche* and *D. bifasciata* because the genes are pseudogenized.

Based on the results of our targeted sequencing, we conclude that all four Arp paralogs were present in the common ancestor of the *obscura* group (Fig. 2, Supplementary Data S3). *Dup1* is present in all surveyed species, whereas *Arp2D* is present in all species except for *D. subobscura*, which has a single base pair deletion leading to a frameshift and numerous stop codons (Supplementary Fig. S4). *Dup2* and *Dup3* have been pseudogenized in the lineage leading to *D. subobscura* and *D. guanche*, while Dup3 has been additionally pseudogenized in the lineage leading to *D. bifasciata* and *D. imaii* (Fig. 2). The published genome sequence of *D. miranda* (Zhou and Bachtrog 2012; Gramates, et al. 2017) had suggested that *Dup2* and *Arp2D* have pseudogenized in this species. However, our survey of 8 strains of *D. miranda* found that all four Arp paralogs are intact in this species (Supplementary Fig. S5, Supplementary Data S2). Overall, our analyses indicate that the Arp paralogs originated ∼14 million years ago in the common ancestor of the *obscura* group of *Drosophila* species, following which only Dup1 has been strictly retained. *D. subobscura* appears unusual in having pseudogenized three of the four Arp *obscura*-specific Arp paralogs whereas the majority of other *obscura* group species have retained all four paralogs.

Intriguingly, even though all the novel Arp paralogs appear to have arisen in the common ancestor of the *obscura* group of *Drosophila* species, they appear to be derived from independent duplication events. Based on initial phylogenetic analyses, we were able to ascribe parentage to only two of the four *obscura*-specific paralogs: Dup1 is 96% identical to an actin protein whereas Arp2D is 70% identical to parental Arp2 in *D. pseudoobscura*. Taking advantage of our additional sequencing within the *obscura* clade (Fig. 2; Supplementary Data S3), we performed additional phylogenetic analyses to delineate the origins of the four Arp paralogs in the *obscura* clade (Supplementary Fig. S6). We find that each paralog forms a monophyletic clade, with relationships that are largely consistent with the branching topology of *obscura* species (Supplementary Fig. S3, S6). Although this analysis still did not have high enough confidence (or bootstrap support) to assign parentage to the Dup2 and Dup3 paralogs (Fig. 1B, Supplementary Fig. S6), we could nonetheless confidently conclude that they represent distinct evolutionary innovations from Dup1.

### The *obscura*-specific Arp paralogs are primarily expressed in males

Publicly available RNA-seq data in *D. pseudoobscura* tissues (Celniker, et al. 2009) confirms that all canonical Arp paralogs are expressed in all tissues. In contrast, all *obscura*-specific Arp paralogs are testis-specific in *D. pseudoobscura* (Celniker, et al. 2009). We investigated whether the male-specific expression of these Arp paralogs observed in *D. pseudoobscura* has been conserved in other species from the *obscura* group. We isolated RNA from males and females of six representative *obscura* species and generated cDNA to conduct RT-PCR analyses, allowing us to compare expression between males and females (Supplementary Data S1). We found that canonical *actin* and *Arp2* are expressed at comparable levels between females and males of each species (Fig. 2, Supplementary Fig. S7). In contrast, all *obscura*-specific Arp paralogs appear to be predominantly expressed in males with low expression detected in females in some species, such as Dup1 in *D. algonquin*. Therefore, the male-specific expression of the *obscura*-specific Arp paralogs is largely conserved since their origin ∼14 mya.

### The *obscura*-specific Arp paralogs have evolved under positive selection

Although Arp genes are typically very highly conserved, testis-specific proteins are often under selective pressure to diversify (Jagadeeshan and Singh 2005; Kleene 2005; Turner, et al. 2008). Therefore, we investigated the selective constraints acting on the male-enriched Arp paralogs present in the *obscura* clade. We first tested each Arp paralog for positive selection using the McDonald-Kreitman test (McDonald and Kreitman 1991), which compares the ratio of nonsynonmous to synonymous fixed differences between two species (D_N_/D_S_) to the ratio of nonsynonmous to synonymous polymorphisms within a species (P_N_/P_S_). If the ratio of fixed differences is far greater than the polymorphism ratio (D_N_/D_S_ ≫ P_N_/P_S_), then this excess of fixed nonsynonymous differences is inferred to be the result of positive selection. We sequenced each of the four Arp paralogs in 10 or 11 *D. pseudoobscura* strains (Supplementary Data S4) and 8 strains of the closely related *D. miranda* (Supplementary Data S2). For each Arp paralog, we aligned the nucleotide sequences and then conducted the MK test, comparing D_N_/D_S_ to P_N_/P_S_ in *D. pseudoobscura* versus *D. miranda* strains. We found that all 4 paralogs have evolved under positive selection with very high statistical significance (Table 1).

**Table 1.** McDonald-Kreitman tests for positive selection. The McDonald-Kreitman compares the ratio of nonsynonmous to synonymous fixed differences between two species (D_N_/D_S_) to the ratio of nonsynonmous to synonymous polymorphisms within a species (P_N_/P_S_). Under neutrality, we expect D_N_/D_S_ ∼ P_N_/P_S_. However, if the ratio of fixed differences is far greater than the polymorphism ratio (D_N_/D_S_ >> P_N_/P_S_), then this excess of fixed nonsynonymous differences (evaluated with a Fisher ‘s exact test) is inferred to be the result of positive selection. Alpha, or neutrality index, represents the proportion of nonsynonymous substitutions likely driven by positive selection (Smith and Eyre-Walker 2002). Alpha is defined as [1-(D_S_P_N_/D_N_P_S_) and is expected to be zero under neutrality, and approaches 1 if all the nonsynonymous substitutions are likely to be driven by positive selection. (Alpha values of 0.8 or higher are considered very strong evidence of positive selection.

| Arp<br>paralog | Polymorphism |  | Divergence |  | # *D. pse*<br>strains | # *D. mir*<br>strains | p-value | Alpha |
| --- | --- | --- | --- | --- | --- | --- | --- | --- |
|  | P<sub>S</sub> | P<sub>N</sub> | D<sub>S</sub> | D<sub>N</sub> |  |  |  |  |
| *Dup1* | 20 | 2 | 8 | 7 | 11 | 8 | **0.008** | **0.885** |
| *Dup2* | 25 | 12 | 4 | 18 | 11 | 8 | **0.000** | **0.893** |
| *Dup3* | 14 | 3 | 6 | 22 | 10 | 8 | **0.000** | **0.941** |
| *Arp2D* | 8 | 8 | 4 | 21 | 10 | 8 | **0.019** | **0.809** |

We next tested for recurrent positive selection at individual sites in the Arp paralogs over the entire *obscura* group using maximum likelihood methods found in the PAML suite (Yang 2007). For each of the four Arp paralogs, all orthologous sequences were aligned based on the codon level. We investigated these alignments for evidence of recombination using GARD analyses (Kosakovsky Pond, et al. 2006); no significant evidence for recombination was observed. Using a species tree, we tested whether NsSites models that permitted codons to evolve under positive selection (M8) were a more likely fit to the data than those models (M7, M8a) that disallowed it. We found marginal evidence for recurrent positive selection acting on *Dup3* but no evidence in the other three paralogs (Table 2). These results contrast with our findings of strong episodic positive selection based on the MK test. Together, they suggest that Arp paralogs have evolved under positive selection, but this positive selection was not driven by recurrent amino acid replacement at a subset of ‘hotspot’ sites; rather, the signal of positive selection appears to be distributed over all four Arp paralogs. This implies that there is selective pressure for gametic Arps to doversify across the actin fold rather than a single surface.

**Table 2.**
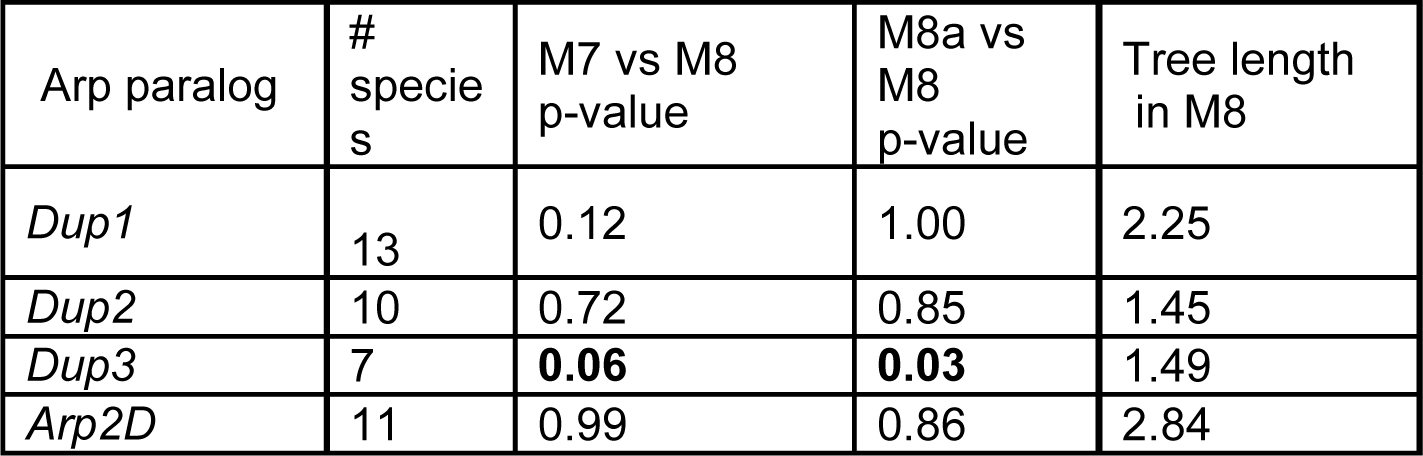
Maximum-likelihood tests for recurrent positive selection. Using the PAML suite (Yang 2007), we tested whether NsSites models that permitted codons to evolve under positive selection (M8) were a more likely fit to the data than those models (M7, M8a) that disallowed it. Tree length refers to the number of nucleotide substitutions per codon, giving an indication of the divergence of the data set. The results we present are from codeml runs using the F3×4 codon frequency model and initial omega 0.4.

| Arp paralog | # species | M7 vs M8 p-value | M8a vs M8 p-value | Tree length in M8 |
| --- | --- | --- | --- | --- |
| *Dup1* | 13 | 0.12 | 1.00 | 2.25 |
| *Dup2* | 10 | 0.72 | 0.85 | 1.45 |
| *Dup3* | 7 | **0.06** | **0.03** | 1.49 |
| *Arp2D* | 11 | 0.99 | 0.86 | 2.84 |

### Evolutionary origins and cytological localization of *Arp2D* in the *obscura* group

We next focused on the evolutionary origins and functional specialization of the more divergent Arp2D. Whereas *Arp2* contains 6 introns, *D. pseudoobscura Arp2D* has none (Fig. 3B, Supplementary Fig. S2). Lack of introns is a conserved feature in all *obscura Arp2D* and leads to a size difference in the genomic PCR analyses based on primers to segments conserved between *Arp2* and *Arp2D* (Fig. 3A). The same PCR reaction using *D. melanogaster* genomic DNA resulted in a single band expected for the size from *Arp2*, verifying the absence of *Arp2D* in this species (Fig. 3A).

**Fig. 3:**
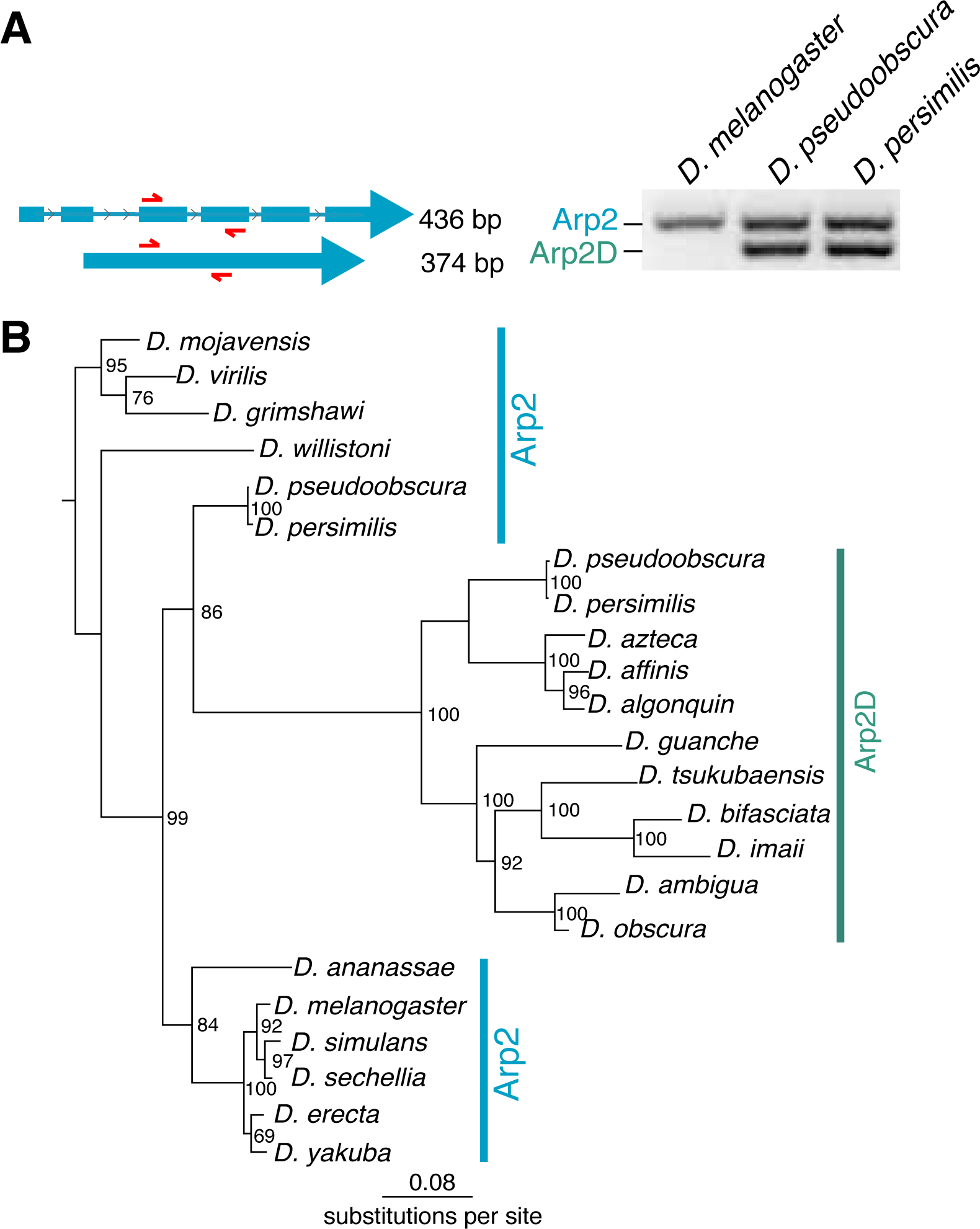
*Arp2D* originated from a retroduplication event of *Arp2*. **A)** Primers targeting sequences in two neighboring exons conserved in *D. pseudoobscura* Arp2 and Arp2D were used to PCR amplify the genomic region in *D. melanogaster, D. pseudoobscura*, and *D. persimilis*. PCR products were run on a 2% agarose gel. Since Arp2D does not contain introns, the PCR product is smaller (374 bp) than that of Arp2 (436 bp). **B)** Nucleotide sequences from Arp2 (from 12 sequenced *Drosophila* species) and Arp2D sequences from the *obscura* group were aligned. A PhyML tree (Guindon, et al. 2010) with 100X resampling was generated and presented using the species from the Drosophila subgenus (*D. virilis, D. mojavensis, D. grimshawi*) as an outgroup.

Using a codon-based alignment, we performed phylogenetic analyses using maximum likelihood methods to investigate the evolutionary origins of *Arp2D* (from 11 *obscura* species) relative to its parental gene *Arp2* (from 12 *Drosophila* species). Our analyses reveal that the *Arp2D* sequences form a monophyletic clade, which arose from within the *Arp2* lineage specifically from the branch that gave rise to *obscura* group species *D. pseudoobscura* and *D. persimilis* Arp2 (86% bootstrap support, Fig. 3B). The branching topology of the *Arp2D* clade (Fig. 3B) also mirrors that of the *obscura* species tree (Fig. 2A) (Barrio and Ayala 1997; Russo 2013). Therefore, we conclude that *Arp2D*, given its lack of introns, arose via retroduplication of the mRNA encoding *Arp2* in the common ancestor of the *obscura* lineage.

### Computational modeling predicts biochemical properties of Arp2D

Next, we investigated whether Arp2D diverged sufficiently to differ in its biochemical properties from Arp2. Arp2 requires incorporation into a multiprotein complex to polymerize branched actin networks (Pollard 2007). This 7-membered complex binds to a pre-formed “mother” actin filament, which leads to a conformational change enabling the complex to bind “daughter” actin monomers; these monomers serve as a platform upon which a second actin filament can polymerize (Pollard 2007). We find that Arp2D, as well as the other Arp paralogs, has preserved the sequence motif required for ATP binding and hydrolysis (Supplementary Fig. S8), which is required for polymerization of actin.

To further compare Arp2D’s biochemical properties to Arp2, we used computational modeling to assess how well Arp2D can sustain the required interactions in the active Arp2-multiprotein assembly structure required to catalyze branched actin networks. We started with a structure of the mammalian Arp2/3 branch junction complex (Pfaendtner, et al. 2012), which was developed by combining data from X-ray structures of the complex along with electron tomography data on the junction conformation (Rouiller, et al. 2008). We used this mammalian model to construct a *D. pseudoobscura* homology model of the entire Arp2/3 complex *i*.*e*., with *D. pseudoobscura* protein sequences for each component of the complex (Arpc1-5, Arp3, and Arp2 or Arp2D) (see Methods). If Arp2D’s sequence were incompatible with Arp2’s canonical structure or function, we would expect to reveal differences in the stability of these complexes upon molecular dynamics (MD) simulation. However, upon comparing the simulations using the *D. pseudoobscura* Arp2/3 and Arp2D/3 complexes including two actin monomers for a nascent branch, we found that both complexes remained intact after a ∼200 ns MD simulation (Fig. 4A, Supplementary Fig. S9A). No steric clashes between residues remained within the structure after the MD. Moreover, the behavior of the daughter actin monomers did not deviate far from the orientation predicted for a stable actin filament (Supplementary Fig. S9A). If the structure were unstable, clashes would be unresolvable by MD simulation and the complex would either fall apart or greatly deform. Instead, our analyses suggest that Arp2D is biochemically capable of replacing Arp2 in the complex.

**Figure 4:**
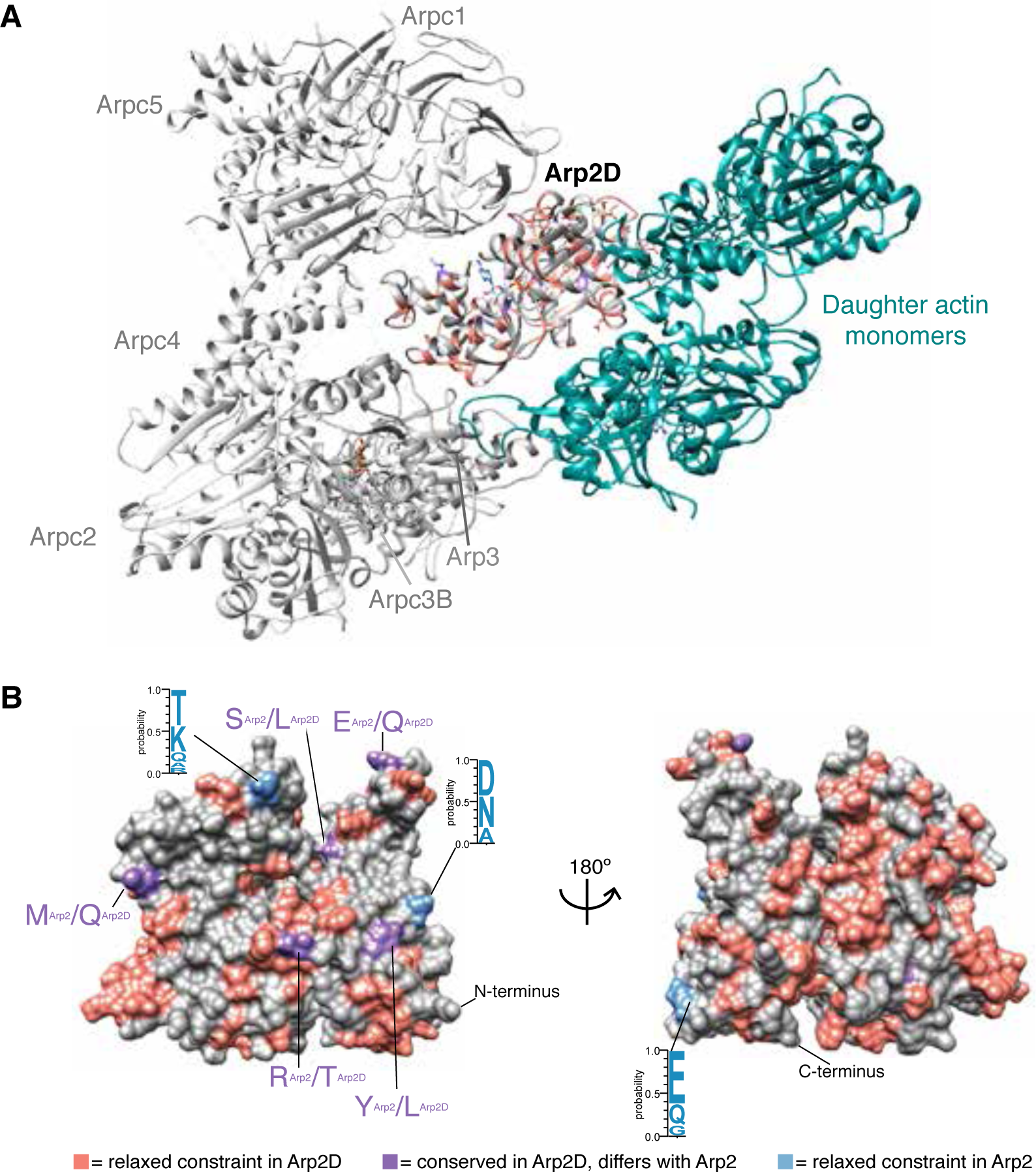
Arp2D is predicted to bind a daughter actin monomer. **A)** A homology model of the *D. pseudoobscura* Arp2/3 complex bound to daughter actin monomers was subjected to structural refinement and equilibration with Arp2D replacing Arp2. Arp2D is color-coded as in (B). Components of the complex are labeled, with daughter actin monomers in teal. **B)** *D. pseudoobscura* Arp2 is shown as a space-filled homology model. All Arp2D protein sequences and a few representative Arp2 sequences (from *D. persimilis, D. pseudoobscura, D. guanche, D. bifasciata* and *D. azteca*) were aligned and residues were categorized as one of the following: 1) relaxed constraint or unconserved in Arp2D (pink), 2) conserved in Arp2D but differs from conserved residues in Arp2 (purple) and 3) unconserved Arp2 residues (blue). Logo plots for residues that are not conserved among Arp2 sequences are displayed.

To further deduce the differences in biochemical properties of Arp2D and parental Arp2, we identified all fixed changes that distinguish *obscura* Arp2D orthologs from all Arp2 orthologs. Based on the alignment of *D. pseudoobscura* Arp2 and Arp2D from all *obscura* species, we classified each residue as a fixed change, relaxed constraint (variable among species) in Arp2D or relaxed constraint in Arp2 (Fig. 4B). We found that there were 9 fixed changes in Arp2D (Fig. 4B, Supplementary Fig. 9B) and only 3 residues that show relaxed constraint in Arp2, predominantly on one surface of Arp2D (Fig. 4B). In contrast, we find that there are many residues under relaxed constraint in Arp2D (Fig. 4B) consistent with our findings of a lower degree of constraint and positive selection of Arp2D. We projected these distinguishing residues onto the surface of Arp2D within the simulated Arp2D/3 complex homology model (Fig. 4A). Notably, we found that the fixed differences between Arp2D and Arp2 made few contacts with other components of the Arp2/3 complex or with the actin monomer (Fig. 4A). Therefore, based on the computational modeling, we infer that Arp2D should be able to catalyze branched actin networks, similar to Arp2.

### Arp2D localizes to gametic actin structures in *D. pseudoobscura* testes

Since Arp2D may have similar biochemical properties as Arp2, we explored whether Arp2D’s cytological localization *in vivo* resembles that of parental Arp2. We first compared Arp2D’s subcellular localization to that of Arp2 in tissue culture cells. Similar to actin, canonical Arp2 localizes at the cell membrane and at cell-cell junctions in tissue culture cells for the polymerization of branched actin networks (Pollard and Borisy 2003). We expressed *Arp2-GFP* in *D. pseudoobscura* cells, and as expected, Arp2 localizes to the cell cortex (Fig. 5A). We then expressed *Arp2D-GFP* and found that it also localizes to the cell cortex and at sites of cell-cell contact (Fig. 5A). This localization, in spite of Arp2D’s ∼30% protein divergence from canonical *D. pseudoobscura* Arp2, suggests that Arp2D either binds directly to actin or it can replace canonical Arp2 in the 7-membered complex that includes Arp3; both scenarios are consistent with our computational modeling analyses (Fig. 4A, Supplementary Fig. S9A).

**Figure 5:**
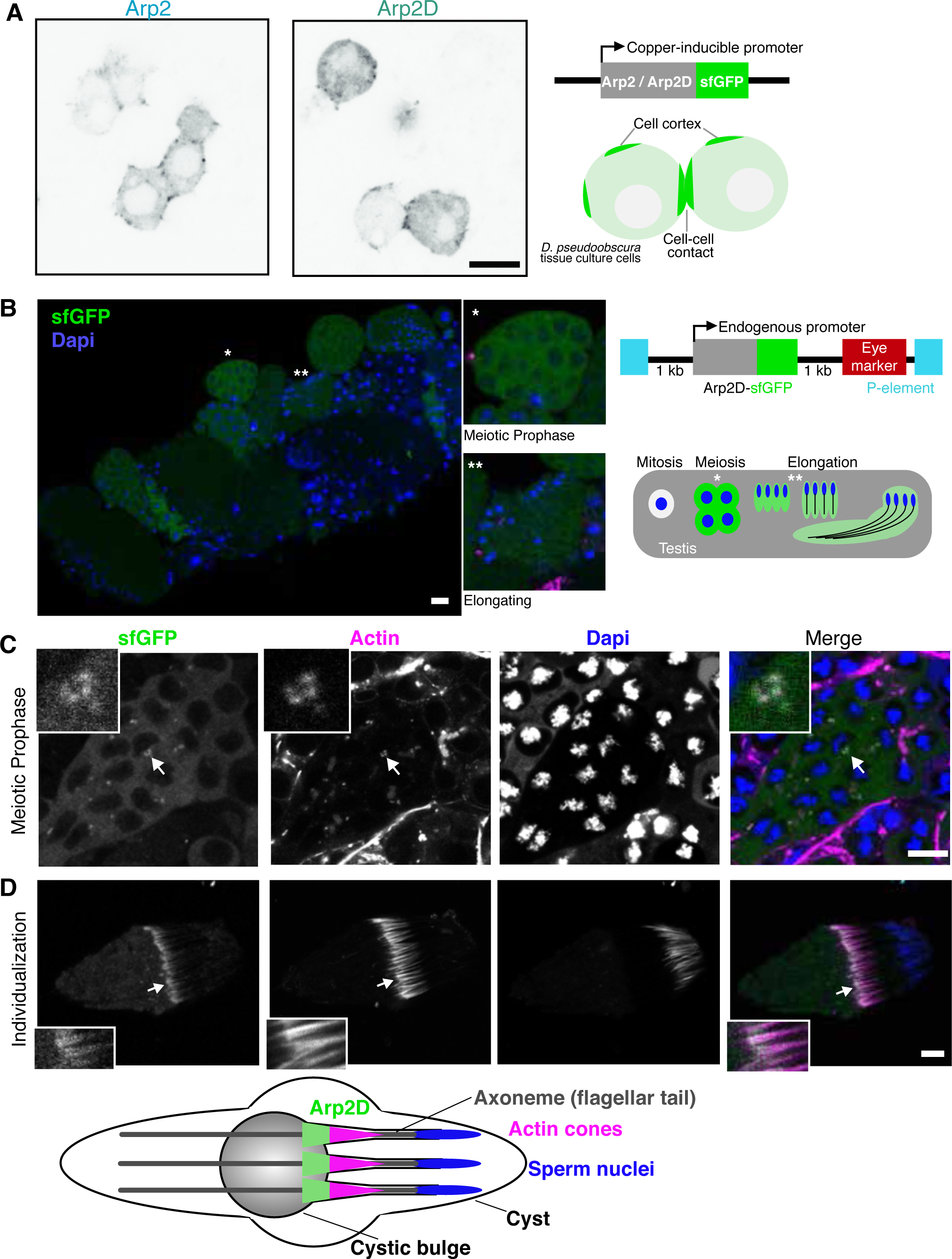
Arp2D is expressed in post-mitotic cell stages and localizes to actin structures. **A)** Arp2 and Arp2D with C-terminal GFP tags were expressed in *D. pseudoobscura* tissue culture cells and plated on concanavalin A-coated plates, followed by live cell microscopy. Scale bar is 7.5 µm. A schematic of the expression construct and the resulting localization are shown. **B)** Cysts at varying stages of spermatogenesis from an *Arp2D-sfGFP D. pseudoobscura* transgenic male. Arp2D is shown in green and DNA in blue. Enlarged images of specific cysts (denoted with asterisks) in stages of meiotic prophase and spermatid elongation are shown. The transgenesis construct injected into *D. pseudoobscura* w^-^ flies is displayed with a schematic indicating the stages of spermatogenesis in which GFP fluorescence is visualized. The number of cells shown at each stage in the schematic are only representative, with 128 spermatids actually following meiosis in *D. pseudoobscura* (Swallow and Wilkinson 2002). **C)** A meiotic cyst, indicated by the nuclear morphology, from the testis of an *Arp2D-sfGFP D. pseudoobscura* transgenic male. Arp2D-sfGFP is green, actin is magenta, and DNA (DAPI stain) is blue. The inset shows GFP concentrated at actin-enriched sites. **D)** A spermatid cyst undergoing individualization with an actin cone enlarged in the inset. Below is a schematic indicating the process of individualization with Arp2D localizing to the front of actin cones (spermatids are representative of 128 per cyst). Scale bars in B-D are 10 µm.

We next assessed Arp2D expression and localization *in vivo*. Using P-element-mediated transgenesis (Thibault, et al. 2004), we generated a transgenic *D. pseudoobscura* line encoding *Arp2D-sfGFP* (superfolder GFP) under the control of *Arp2D*’s native promoter (Fig. 5B). We placed sfGFP specifically at the C-terminus of Arp2D since Arp2 has been shown to be functionally unperturbed with a C-terminal tag (Egile, et al. 2005). We confirmed the expression of the full-length protein by western blot analysis and found that the Arp2D-sfGFP protein was expressed at the expected size (Supplementary Fig. S10). Consistent with the RNA-seq and RT-PCR analyses, we found that this transgene was expressed in testes. To image Arp2D localization *in vivo*, we dissected testes and imaged live GFP fluorescence with the addition of live-imaging probes specific for DNA and actin (Fig. 5B). By confocal microscopy, we found that Arp2D-sfGFP is clearly expressed in meiotic and post-meiotic stages of spermatogenesis where it localizes to the cytoplasm (Fig. 5B). We confirmed that the fluorescence was specific to sfGFP and not due to autofluorescence by imaging *D. pseudoobscura white-* flies lacking *Arp2D-sfGFP* with the same confocal laser settings (Supplementary Fig. S11).

We first detected expression of Arp2D-sfGFP during meiotic prophase, during which Arp2D-sfGFP is often detected at points of concentrated actin at the cell surface that we predict to be endocytic sites (Fig. 5C) (Pollard 2007). Arp2D-sfGFP expression persists throughout sperm individualization. During spermatid elongation following meiosis, Arp2D-sfGFP appears diffuse, albeit at a lower fluorescence intensity than meiotic cells (Fig. 5B). Then Arp2D-sfGFP exhibits striking localization at actin during the final step of spermatogenesis–individualization (Fig. 5D). Since cells are interconnected throughout mitotic and meiotic divisions in spermatogenesis, individualization must take place to separate spermatids. During this process in *D. pseudoobscura*, 128 spermatids in a single cyst (Swallow and Wilkinson 2002) are each separated by an actin cone, which forms at the sperm nucleus and then translocates along the sperm tail, disposing of excess cytoplasm and encasing each spermatid in its own membrane (Fig. 5D) (Noguchi, et al. 2008). Arp2D clearly localizes to motile actin cones (Fig. 5D), which translocate along sperm tails to remove excess cytoplasm and encase each spermatid in its own membrane.

This localization of Arp2D-sfGFP to actin cones is highly reminiscent of the localization of parental Arp2 in *D. melanogaster* (Noguchi, et al. 2008). Arp2 generates branched actin networks towards the front of actin cones, creating a fan-like structure, and this polymerization of branched actin facilitates the motility of the cones (Noguchi, et al. 2008). We find Arp2D localizes to fan-like actin cones that are readily motile and have translocated away from sperm nuclei (Fig. 5D; Supplementary Fig. S12). However, we do not observe Arp2D in immotile cones that are not fan-like (i.e. branched actin networks are absent) (Supplementary Fig. S12). Thus, Arp2D localizes to a subset of previously-described gametic Arp2-containing actin structures *in D. pseudoobscura* testes. Based on its predicted ability to participate in Arp2-related interactions, its similar localization prior to and during spermatogenesis, and its positive selection, we hypothesize that Arp2D allows for the specialization and recurrent innovation of Arp2-related functions specifically in the male germline, without injury to Arp2’s many highly conserved functions in the soma.

## Discussion

In this study, we report a burst of genetic innovation in cytoskeletal genes in one lineage of *Drosophila* species. We find that four Arp paralogs arose independently in the *obscura* group of *Drosophila*. Given the extreme conservation during eukaryote evolution of the cytoskeletal apparatus, particularly of Arp genes, such a burst of genetic innovation is unexpected.

All the *obscura*-specific Arp paralogs are male-biased in expression. This is consistent with the ‘out-of-testis’ hypothesis, which posits that evolutionarily young genes often originate with testis-restricted expression in multiple species before they expand their expression profile to other tissues (Vinckenbosch, et al. 2006; Kaessmann 2010; Assis and Bachtrog 2013). This testis-biased ‘nursery ‘ of young genes is thought to occur for two reasons. First, testes often have inherently more promiscuous transcription allowing duplicate genes a higher likelihood of being transcribed and translated (Vinckenbosch, et al. 2006). Second, testis-expressed genes are also subject to extremely high rates of evolutionary innovation because they often serve as battlegrounds for multiple genetic conflicts (Kleene 2005). As such, evolutionary innovations are more likely to be valuable for testis-specific expression than in other tissues (Vinckenbosch, et al. 2006; Kaessmann 2010). A corollary to this second point is that evolutionary innovation may be detrimental in other tissues especially if it upsets dosage of proteins in multimeric complexes (Holland and Johnson 2018). Given that all the canonical Arp paralogs are ubiquitously expressed, testis-specific innovation of Arp paralogs affords the opportunity for cytoskeletal innovation to occur in the testis without interfering with their essential functions in the soma. Thus, Arp specific duplications might relieve some of the antagonistic pleiotropy between Arp innovation for testis functions and stasis of Arp functions for somatic tissues (Gallach and Betran 2011).

Are the Arp paralogs simply a means to maintain Arp function in the male germline? We can only ascribe parentage to two of the four *obscura*-specific Arp paralogs. In both cases, the parental genes are present on the X chromosome. For instance, *Dup1* is an autosomal duplicate of an X chromosomal *actin* gene, whereas *Arp2D* is a retroposed copy of X-chromosomal *Arp2*. Many X chromosomal genes are subject to meiotic silencing during male meiosis, and it has been previously proposed that this requirement for male germline expression has driven the disproportionate trafficking or retroduplication of several X-chromosomal genes onto autosomes to maintain germline function in both *Drosophila* and mammals (Betran, et al. 2002; Emerson, et al. 2004; Dai, et al. 2006; Vibranovski, Lopes, et al. 2009; Vibranovski, Zhang, et al. 2009). RNA-seq data suggest that canonical Arp2 is indeed expressed in the testis, albeit at lower levels (Celniker, et al. 2009). Under this scenario, however, one might expect to primarily see signatures of preservation rather than divergence of function. Indeed, a recent report showed that maintenance of ancestral functions is more likely to be associated with purifying selection, whereas novel functions are more likely to be associated with positive selection (Jiang and Assis 2017). Our finding that all four Arp paralogs are subject to positive selection, as well as the high divergence of three of the four paralogs from canonical Arps, suggests that it is unlikely that only preservation of function can sufficiently explain this burst of innovation in the *obscura* group. Instead, we propose that duplication of X chromosomal copies both protected and allowed for innovation of male germline-specific expression and function.

What novel function might Arp2D play in the testis? Despite Arp2D’s divergence, fixed residue differences between all Arp2D orthologs and canonical Arp2 lie outside the predicted binding interfaces with actin monomers and the remaining components of the multiprotein complex. Furthermore, our analyses reveal that Arp2D retains the ability to localize to actin similarly to Arp2. Nevertheless, residue differences between Arp2D and Arp2 (Fig. 4B) may alter important protein-protein interactions. Canonical Arp2 requires regulatory proteins to activate its catalysis of branched actin networks; inhibitors of Arp2 have been identified as well (Pollard 2007; Gandhi, et al. 2010). Arp2D’s divergence and rapid evolution might have enabled it to lose these interactions and gain novel ones, which subsequently may affect its catalytic ability to polymerize actin filaments. Any alteration of the rate of actin polymerization would have significant consequences. For example, the motility of actin cones is thought to be generated by the dynamics of Arp2/3-generated actin polymerization (Noguchi and Miller 2003). We hypothesize that divergence of Arp2D function may facilitate a novel meiotic and post-meiotic specialization that may not be possible for Arp2, with its many roles in all tissues beyond the testis.

One of the most remarkable aspects of our study is the finding of four independently-derived Arp paralogs in the same lineage of *Drosophila*. Even if we were to suppose that the pressures of functional novelty or preservation of male germline-specific expression drove this innovation, there is no *a priori* reason why this remarkable pattern of convergence would be expected to all occur only in one lineage of *Drosophila*, especially when one considers that the parental genes for each *obscura*-specific Arp paralog could be distinct. Since the likelihood of the generation of new Arp paralogs is not unique to *obscura*, we wondered if there is an aspect of spermatogenesis in the *obscura* group that uniquely influences the retention of the Arp paralogs.

Among investigated *Drosophila* species, the *obscura* group is unique in possessing sperm heteromorphism: the simultaneous production of distinguishable types of sperm by a single male (Joly 1989, 1991; Joly 1994; Swallow and Wilkinson 2002; Holman, et al. 2008). For example, *D. pseudoobscura* sperm are heteromorphic with fertilization-competent eusperm and two types of parasperm, which cannot fertilize but rather may increase the competitive ability of eusperm to fertilize females (Snook, et al. 1994; Holman, et al. 2008; Holman and Snook 2008; Alpern, et al. 2019). It is unclear how parasperm increase eusperm competitiveness, but proposed mechanisms include displacing rival fertilizing sperm in the female, protection from spermicide in the female reproductive tract and manipulation of female receptiveness to remating (Oppliger 1998; Cook 1999; Holman and Snook 2006, 2008; Alpern, et al. 2019).

Eusperm are easily identified by microscopy because they are ∼5 times longer than parasperm (Holman, et al. 2008; Alpern, et al. 2019). Thus, if sperm heteromorphism is an adaptive trait specific to the *obscura* group of *Drosophila*, it is possible that the dual pressures to produce both eusperm and parasperm may have required significant innovation in the cytoskeletal machinery required to produce the different types of mature sperm through individualization.

Intriguingly, the magnitude of sperm heteromorphism is not static in *obscura* group species, which show significant variation in heteromorphic sperm length and size differences (Holman, et al. 2008). This variance may explain the still ongoing positive selection and genetic turnover of the Arp paralogs in the *obscura* group. In this regard, it may be especially interesting to re-examine sperm heteromorphism in species like *D. subobscura* that lack three of the four *obscura*-specific Arp paralogs or in *D. pseudoobscura* strains in which the Arp paralogs have been experimentally knocked out. Given its rarity, sperm heteromorphism is unlikely to be a universal explanation for the function of testis-specific ‘orphan’ Arp genes in *Drosophila* and mammals but may nevertheless illustrate how male germline specific functions invoke or facilitate lineage-specific adaptation of cytoskeletal functions.

## Methods

### *Drosophila* Species and Strains

Species in the *obscura* clade were either obtained from the National *Drosophila* Species Stock Center (Cornell University) and were a kind gift from Dr. Masayoshi Watada (Ehime University). The *D. pseudoobscura* and *D. miranda strains* were generously provided by Dr. Nitin Phadnis (University of Utah) and Dr. Doris Bachtrog (University of California, Berkeley) respectively. For a list of strains used in this study, see Supplementary Data S1.

### Discovery of Arp duplications in sequenced genomes

The protein sequence of each *D. melanogaster* canonical Arp was used in a tBLASTn search (Altschul, et al. 1997) for other Arp sequences in the 12 sequenced and annotated species of *Drosophila* (Drosophila 12 Genomes 2007; Gramates, et al. 2017). The sequences were identified as canonical Arps based on phylogenetic grouping. The outliers, or the Arp paralogs, were further confirmed as novel Arps by verifying their absence in the syntenic loci of the other sequenced *Drosophila* species. A 10-20 Kb region of the syntenic locus from each *Drosophila* species was collected and a tBLASTn search was conducted against the locus with each paralog’s protein sequence from *D. pseudoobscura* (Kearse, et al. 2012). Of the 12 sequenced and annotated Drosophila species, only *D. persimilis* and *D. pseudoobscura* resulted in positive hits, while the other species were negative.

### Sequencing shared syntenic loci of Arp paralogs in the *obscura* group

Whole flies (10-15) with approximately equal number of males and females were ground in the following buffer: 10 mM Tris pH 7.5, 10 mM EDTA, 100 mM NaCl, 0.5% SDS and 0.5 µg/µL Proteinase K (NEB). The flies were then incubated at 55°C for 2 hrs, followed by a phenol-cholorform extraction. The final ethanol-washed DNA pellet was resuspended in distilled water. To obtain the sequences of Arp paralogs from non-sequenced *obscura* group species, we aligned the syntenic loci from *D. pseudoobscura* and *D. melanogaster* (and for Arp2D, *D. ananassae* was included) and designed primers in conserved intergenic regions or neighboring genes. If gene products were >5 kb, then two PCRs were conducted with forward and reverse primers aligned with the highly conserved ATP-binding motif, which was later re-sequenced after the 5’ and 3’ prime halves of each paralog were obtained. Primers were iteratively designed based on successful PCRs from divergent species. For a full list of primers used, see Supplementary Data S1.

A touchdown PCR protocol was followed (Korbie and Mattick 2008) and Phusion was used per the manufacturer’s instructions (NEB). PCR products that were approximately less than 3kb were TOPO cloned into the pCR4-Blunt vector (Thermo Fisher Scientific) and subsequently sequenced with the M13F and M13R primers. PCR products that were greater than 3kb were directly sequenced. Coding sequences and sequencing of an extended region of each locus can be found in Supplementary Data 3, respectively. Sequences are to in the process of being deposited in Genbank. To differentiate between the presence and absence of *Arp2* and *Arp2D* (Fig. 3A), we designed primers that aligned completely with both genes and flanked an intron in *Arp2* (For: TGATGGTCGGCGATGAGGC, Rev: CGTCAAATGTGGCAGGGCA).

### Sequence Alignments and Phylogenetic Analyses

Protein sequences were aligned with MAFFT (Katoh and Standley 2013) and genes were aligned using translation align in the Geneious software package (version 9.1.3, (Kearse, et al. 2012)). Gaps were removed where less than 80% of the sequences aligned. Maximum likelihood trees were generated using the LG substitution model in PhyML (Guindon, et al. 2010) and analyzed for statistical support using 100 bootstrap replicates. Trees were visualized with Geneious.

### Imaging Arp2D in tissue culture cells

*D. pseudoobscura* tissue culture cells (cell line #ML83-63) were obtained from the *Drosophila* Genomics Resource Center (https://dgrc.bio.indiana.edu/Home) and cultured in M3+BPYE media supplemented with 10% fetal calf serum. Cells were incubated at 25°C and passaged using a cell scraper (Fisher Scientific, 08-771-1A) to facilitate transfers. Cells were transfected with 1 µg DNA and 8 µl Fugene HD (Promega) in serum-free media (100 µL total volume), followed by a 5 min incubation at room temperature. The transfection solution was added drop-wise to a confluent layer of *D. pseudoobscura* cells in a single well of a 6-well dish. The cells were incubated for 24 hours at 25°C and then exogenous gene expression was induced with 100 µM copper every 24 hours and imaged 24 hrs-48 hrs following transfection when a sufficient number of cells displayed fluorescence. Cells were then resuspended and ∼30 µL was added to the coverslip of a MatTek dish (MatTek corporation, P35G-1.5-10-C), coated with 0.5 mg/mL concanavalin A (MP Biomedicals). The media was exchanged with PBS to decrease background when imaging, and cells were imaged using a confocal microscope (Leica TCS SP5 II) and LASAF software (Leica).

### Comparison of Sex-Biased Expression

Six species were chosen that spanned a wide evolutionary distance in the *obscura* group to compare male versus female expression. Whole male and female flies were collected separately and RNA was extracted using TRIzol and further purified per the manufacturer’s instructions (Invitrogen). RNA was then treated with DNAse and used to synthesize cDNA with SuperScript III (Invitrogen). For each reaction, a corresponding reaction without RT was conducted to detect genomic DNA contamination (Supplementary Fig. S7). Subsequent cDNA was then utilized for RT-PCR of ribosomal *Rp49* to assess relative amounts of cDNA among samples. For RT-PCR of the Arp paralogs, primers were designed to yield a ∼100-350 bp product for efficient amplification and to distinguish the paralog from actin or Arp2. For a full list of primers used, see Supplementary Data S1. RT-PCRs were conducted using Phusion per the manufacturer’s instructions (NEB) for 25 cycles. Equal volumes of each RT-PCR were loaded on a 1% agarose gel for analysis.

### Structural Analysis

To analyze conservation and divergence of the proteins, sequences were aligned with MAFFT (Katoh and Standley 2013) and mapped onto the structure’s surface using Chimera. Phyre2 (Kelley, et al. 2015) was used to obtain a homology model of *D. pseudoobscura* Arp2. This software used crystal structures 1O1F and 4JD2, respectively, to model the *D. pseudoobscura* proteins. All structural analysis was done using Chimera or MacPymol as indicated in the figure legends. To analyze Arp2D in the context of the Arp2/3 complex, sequences of all the *D. pseudoobscura* Arp2/3 subunits (Arpc1-5, Arp2D and Arp3) were collected and submitted to Swiss-Model (Waterhouse, et al. 2018) to generate a homology model using an X-ray crystal structure of the *Bos taurus* Arp2/3 complex as a template (PDB 3DXM) (Nolen, et al. 2009). The Arp2D homology model substituted Arp2 in the complex. *D*.*pseudoobscura* actin was modeled using template PDB 2ZWH (Oda, et al. 2009). To view Arp2D with respect to actin, an activated mammalian Arp2/3 junction model was constructed, as described in Supplementary Information.

### Generation of an *Arp2D-sfGFP D. pseudoobscura* transgenic line

The *Arp2D* paralog was encoded with a superfolder GFP (sfGFP) tag and cloned into the restriction sites BamHI and NotI in the pCasper4 vector. Arp2 has shown to be best tagged at the C-terminus; thus a similar strategy was taken with Arp2D. The 1 kb intergenic regions that were upstream and downstream of the paralogs were included to allow for endogenous expression under the control of proximal transcriptional regulatory regions. The constructs were maxi-prepped (Machery-Nagel) for high quality DNA and used for injections of *D. pseudoobscura* w-flies (gift from Nitin Phadnis, University of Utah). Fly embryos were injected by Rainbow Transgenic Flies, Inc. in combination with transposase *in trans*. We crossed larvae that survived the injections to *D. pseudoobscura* w-flies (4:1 or 5:2 females to males) and selected progeny with pigmented eyes, which ranged from light orange to deep red. Transformants that appeared in separate crosses were designated as different founder lines and two founders were identified. Homozygous lines were generated and imaged live.

### Imaging of *Arp2D-GFP* transgenic flies

For live imaging, the testes from male transformants were dissected into phosphate-buffered saline (PBS) and then transferred to a drop of PBS on a slide. PBS was exchanged for PBS containing the DNA stain Hoechst and sir-actin (10 µM; Cytoskeleton, Inc), which is used for live samples. The testes were then torn open with tweezers to release all cell-types during spermatogenesis; this technique allowed for improved visibility of different subpopulations of developing sperm. The tissue was incubated in the imaging solution for 5 minutes in the dark and then a coverslip was placed on top. The sample was immediately imaged using a confocal microscope (Leica TCS SP5 II) and LASAF software (Leica) for 20-30 min, until GFP signal faded and signs of apoptosis were visible. We opted for live imaging of the transformants to avoid the possibility of fixation artifacts; we also found immunofluorescence of the samples led to a high background, making it difficult to know what was true sfGFP fluorescence.

### Analysis of Positive Selection

To test for positive selection at the population level, we compared sequences of all four Arp paralogs in 10-11 strains of *D. pseudoobscura* to those in 8 strains of *D. miranda*. The Arp paralogs were sequenced with primers that aligned within the 5’ and 3’ UTRs of the *D. pseudoobscura* genes. The genes were codon-based aligned and subjected to the MK test with an online resource (Egea, et al. 2008). To test for site-specific positive selection, we used the PAML suite (Yang 2007). The coding sequences for each paralog from all *obscura* species were codon-based aligned using Geneious (Kearse, et al. 2012) and all gaps were removed. The requisite format of the alignments was obtained using Pal2Nal (Suyama, et al. 2006). The alignment and corresponding species tree for each paralog were then used as input files for the program CODEML NSites in the PAML suite. The starting omega used was 0.4 with a codon frequency model of F3×4. Tests with different starting omegas and codon frequency models yielded similar results. The models M7, M8a and M8 were compared to determine if positive selection was present. The test indicated whether the evolution of the paralogs fit a model that allows for DN/DS > 1 (M8) or models that do not allow for D_N_/D_S_ >1 (M7 and M8a). We used the M8 Bayes Empirical Bayes analysis to identify specific residues under positive selection.

## Acknowledgments

We thank Lisa Kursel, Antoine Molaro and Jeannette Tenthorey for their comments on the manuscript, as well as other members of the Malik lab for helpful discussions. We also greatly appreciate the gift of fly strains from Nitin Phadnis (University of Utah), Doris Bachtrog (University of California Berkeley), the National *Drosophila* Species Stock Center (Cornell University), Masayoshi Watada (Ehime University). Transgenic *D. pseudoobscura* flies were generated by Rainbow Transgenic Flies, Inc. Molecular dynamics simulations were performed on resources provided by NYU IT High Performance Computing. This work was funded by the Jane Coffin Childs Memorial Fund (CMS), startup funds from NYU (GMH), NIH grant R01GM074108 (HSM) and the Howard Hughes Medical Institute (HSM). HSM is an Investigator of the Howard Hughes Medical Institute.

## Supplementary Information

### Supplementary Figure Legends

**Supplementary Figure S1: Sequenced *Drosophila* species encode canonical Arps and lineage-specific Arps**.

**A)** All surveyed *Drosophila* species have a representative from each subfamily of canonical Arps, and the genomes of only *D. pseudoobscura* and *D. persimilis* encode Dup1-3 and Arp2D. **B)** The duplicates are encoded in different chromosomal locations in *D. pseudoobscura* than canonical actins, muscle actins, and Arp2. **C)** The syntenic locations of the duplicates found in *D. pseudoobscura* and *D. persimilis* do not encode an actin-like gene in the other sequenced and annotated *Drosophila* species. Genes are named based on the *D. melanogaster* names, and the colors differentiate genes used to define the syntenic locus. Gray arrows (without names) do not define the locus, as they are not present in most of the species, and a gray box outlines the loci in which the duplicates are present.

**Supplementary Figure S2: Arp2D is present in only the *obscura* group**.

The syntenic locus of Arp2D and Arp2 in the 12 sequenced *Drosophila* species is displayed. The approximate exon and intron gene models of Arp2 is shown.

**Supplementary Figure S3: Nucleotide trees including pseudogenized genes recapitulate overall species tree topology**.

For each Arp duplicate, coding ORFs were codon-based aligned and then consensus aligned with pseudogenized genes, which were not included in Figure 4A. Maximum-likelihood trees with 100X resampling were generated with PhyML (Guindon, et al. 2010).

**Supplementary Figure S4: A single base-pair deletion leads to pseudogenization of Arp2D in *D. subobscura***.

The Arp2D sequences from *D. pseudoobscura* and *D. subobscura* are aligned and a single base pair deletion in *D. subobscura* is indicated with a blue box with early stop codons noted with red boxes.

**Supplementary Figure S5: *D. miranda* encodes full-length Dup2 and Arp2D**.

**A)** The sequence of Dup2 from one *D. miranda* strain is aligned with the corresponding nucleotide sequence on Flybase (Attrill et al., 2016). The blue box indicates a gap in the sequence from Flybase and the red boxes indicate the resulting stop codons. **B)** The Arp2D sequence from one *D. miranda* strain is aligned with the corresponding nucleotide sequence from Flybase and two sites within the sequence from Flybase exhibit gaps resulting in numerous stop codons.

**Supplementary Figure S6: Gene and protein trees of Duplicates 1-3 and canonical Arps**

**A)** Gene tree of actin (from 12 sequenced *Drosophila* species) and duplicates 1-3 from *obscura* species. Sequences of Arp2-10 from *D. melanogaster* and *D. pseudoobscura* were included. Translation aligned and PhyML tree with 100X resampling (Guindon, et al. 2010). Gaps with 80% or more unaligned sequences were removed. **B)** Protein sequences from those represented in (A) were aligned with MAFFT and a PhyML tree with 100X resampling is shown. High bootstrap indicates Dup1 is a duplicate of actin. Parentage of Dup2 and Dup3 is unclear in (A) and (B) due to low bootstrap support.

**Supplementary Figure S7: RT-PCR indicates male-enriched expression of the Arp duplicates**.

RT-PCR was conducted with representative species in the *obscura* clade to compare expression of the Arp duplicates (Dup1, 2, 3, and Arp2D), canonical actin (Act5C) and canonical Arp2 between males and females. PCR for *Rp49* was also conducted on cDNA samples generated with (+) and without (-) reverse transcriptase to indicate lack of genomic DNA contamination in the samples. PCR of Rp49 also revealed similar levels of sample loaded for the sexes of each species. The asterisk denotes the band of interest to distinguish it from non-specific bands.

**Supplementary Figure S8: The ATP-binding motifs are conserved in the *D. pseudoobscura* Arp duplicates**.

Alignments of the ATP-binding motifs in canonical actin (human ActB and *D*.*pse* Act5C), Arp2, and the *D. pse* Arp duplicates are shown. Motifs 1 and 2 are important for binding the phosphates of ATP and motif 3 is important for interacting the nucleoside. Residues within a red box have been shown in the actin structures to bind the ATP ligand. Residues with an asterisk are invariant and shown to be critical catalytic residues. The denoted Asp in particular coordinates the divalent metal ion required for phosphor-transfer.

**Supplementary Figure S9: Arp2D/3 exhibits a stable complex similar to Arp2/3**.

**A)** Graphs indicate the stability of the models with a daughter actin filament. The top graph shows the C_α_ RMSD of the *D*.*pse* complex (Arp2 or Arp2D, Arp3, RPC1-5) relative to the initial input model. The steady RMSD value following 100 ns of dynamic simulation suggests the structure of the complex is stable. The bottom two graphs show the daughter actin monomer ‘s subdomain 2-1-3-4 dihedral angle, as defined in (Saunders and Voth 2012; Hocky, et al. 2016). Since the *D*.*pse* Arp2D/Arp3 complex exhibits an RMSD and dihedral angle similar to *D*.*pse* Arp2/3, it is behaving biochemically like canonical Arp2 and can most likely adopts the conformation required for generating a daughter actin filament. **B)** Fixed changes between Arp2 and Arp2D are listed. The Arp2D residues are conserved within all *obscura* species.

**Supplementary Figure S10: Full-length Arp2D-sfGFP protein is expressed in the testis**.

Thirty testes were dissected from two founder lines (“line 7 and 9”) encoding Arp2D-sfGFP and were denatured using SDS loading buffer. Thirty-two testes were dissected from a negative control, a *D*.*pseudoobscura* transenic encoding the mini-white gene. Top image: probed with anti-GFP. The purple arrow denotes the band of expected size for Arp2D-sfGFP, 73 kDa. Asterisks indicate non-specific bands. Two different volumes (25% and 75% of the total sample) were loaded for the two fly lines. Bottom: probed with anti-tubulin, expected size is 50 kDa.

**Supplementary Figure S11: *D. pseudoobscura* w-mature sperm exhibit autofluorescence but none in meiotic and post-meiotic cysts**.

Cysts at different stages of spermatogenesis from a *D. pseudoobscura* w^-^ male fly were imaged with the same laser intensity settings for GFP fluorescence as with the Arp2D-sfGFP *D. pseudoobscura* transgenic. Top row displays elongating spermatids and a mature cyst of eusperm. Second row displays meiotic cells and elongating spermatids. GFP is in green and DNA is in blue. Scale bars are 20 µm.

**Supplementary Figure S12: Arp2D localizes only to motile fan-like actin cones**

Top panel shows a cyst undergoing developing actin cones that are not yet motile for individualization to take place. Bottom panel displays a cyst with motile actin cones that are fan-like in shape due to branched actin networks. Arp2D is green, actin is magenta and DNA is blue. Scale bars are 10 µm.

### Supplementary Data

**Supplementary Data S1: Primers used in study**

**Supplementary Data S2: Arp duplicate sequences in *D. miranda* strains**

**Supplementary Data S3: *Obscura* species ‘ Arp duplicate coding sequences and loci**

**Supplementary Data S4: Arp duplicate sequences in *D. pseudoobscura* strains**

**Supplementary Methods text for Arp2/3 complex model and molecular dynamics simulations**

The initial all-atom model of an activated Arp2/3 complex was built following the protocol of (Pfaendtner, et al. 2012), but using a newer model for the actin filament, namely that of (Oda, et al. 2009), PDB ID 2ZWH. A single structure from simulations of a full junction by one of us (GMH) is shown in (Aydin, et al. 2018). In brief, to build the junction model, actin monomers in the filament had a magnesium ion and a bound ADP as well as coordinating waters placed inside, and then assembled into a filament as previously described (Saunders and Voth 2012; Hocky, et al. 2016). A mother filament of 13 subunits and a daughter filament of 11 subunits in ideal geometry were constructed. These filaments were aligned to the Branch10 structure in (Pfaendtner, et al. 2012) by C_α_ RMSD. Mother actin subunits 7 and 9 were replaced with structures from (Pfaendtner, et al. 2012) because these are in direct contact with the complex, and come from (Rouiller, et al. 2008)’s reconstruction but the coordinating waters from the equilibrated Oda structure are placed into these two subunits by alignment of the actin subunits. The Arp2/3 complex structure from Branch10 in (Pfaendtner, et al. 2012) was preserved, except that a magnesium ion, water, and an ADP molecule were placed inside of Arp2, and the same was done for Arp3 except with an ATP molecule/water/ion from an equilibrated ATP-bound Oda monomer (Katkar, et al. 2018) based on the structure in PDB ID 1J6Z, due to the difference in nucleotide hydrolysis rates of Arp2 and Arp3 (Pollard 2007). In the case of this work, this whole protocol was followed to build a mammalian junction, but only the first two actin subunits in the daughter filament were kept. The system was then solvated in TIP3P water with 0.18M KCl. Equilibration was performed in NAMD (Phillips, et al. 2005) using the CHARMM22+CMAP forcefield following the exact procedure in (Hocky, et al. 2016). This structure formed the basis for the homology models of the Arp2/3 complex and daughter actin proteins used in this work. Subsequently, the homology models of the complex and actin were re-aligned with this structure and equilibrated according to this same protocol using NAMD. Then subsequent MD simulations were performed in GROMACS (Abraham, et al. 2015).

## References

Abraham MJ, Murtola T, Schulz R, Szilard P, Smith JC, Hess B, Lindahl E. 2015. GROMACS: High performance molecular simulations through multi-level parallelism from laptops to supercomputers. SoftwareX 1-2:19–25.

Alpern JHM, Asselin MM, Moehring AJ. 2019. Identification of a novel sperm class and its role in fertilization in Drosophila. J Evol Biol 32:259–266.

Altschul SF, Madden TL, Schaffer AA, Zhang J, Zhang Z, Miller W, Lipman DJ. 1997. Gapped BLAST and PSI-BLAST: a new generation of protein database search programs. Nucleic Acids Res 25:3389–3402.

Assis R, Bachtrog D. 2013. Neofunctionalization of young duplicate genes in Drosophila. Proc Natl Acad Sci U S A 110:17409–17414.

Aydin F, Katkar HH, Voth GA. 2018. Multiscale simulation of actin filaments and actin-associated proteins. Biophys Rev 10:1521–1535.

Babcock CS, Anderson WW. 1996. Molecular evolution of the Sex-Ratio inversion complex in Drosophila pseudoobscura: analysis of the Esterase-5 gene region. Mol Biol Evol 13:297–308.

Barrio E, Ayala FJ. 1997. Evolution of the Drosophila obscura species group inferred from the Gpdh and Sod genes. Mol Phylogenet Evol 7:79–93.

Betran E, Thornton K, Long M. 2002. Retroposed new genes out of the X in Drosophila. Genome Res 12:1854–1859.

Blessing CA, Ugrinova GT, Goodson HV. 2004. Actin and ARPs: action in the nucleus. Trends Cell Biol 14:435–442.

Boeda B, Knowles PP, Briggs DC, Murray-Rust J, Soriano E, Garvalov BK, McDonald NQ, Way M. 2011. Molecular recognition of the Tes LIM2-3 domains by the actin-related protein Arp7A. J Biol Chem 286:11543–11554.

Cairns BR, Erdjument-Bromage H, Tempst P, Winston F, Kornberg RD. 1998. Two actin-related proteins are shared functional components of the chromatin-remodeling complexes RSC and SWI/SNF. Mol Cell 2:639–651.

Celniker SE, Dillon LA, Gerstein MB, Gunsalus KC, Henikoff S, Karpen GH, Kellis M, Lai EC, Lieb JD, MacAlpine DM, et al. 2009. Unlocking the secrets of the genome. Nature 459:927–930.

Cook PA, Wedell, N. 1999. Non-fertile sperm delay female remating. Nature 397:486–486.

Dai H, Yoshimatsu TF, Long M. 2006. Retrogene movement within- and between-chromosomes in the evolution of Drosophila genomes. Gene 385:96–102.

Dominguez R, Holmes KC. 2011. Actin structure and function. Annu Rev Biophys 40:169–186.

Drosophila 12 Genomes C. 2007. Evolution of genes and genomes on the Drosophila phylogeny. Nature 450:203–218.

Egea R, Casillas S, Barbadilla A. 2008. Standard and generalized McDonald-Kreitman test: a website to detect selection by comparing different classes of DNA sites. Nucleic Acids Res 36:W157–162.

Egile C, Rouiller I, Xu XP, Volkmann N, Li R, Hanein D. 2005. Mechanism of filament nucleation and branch stability revealed by the structure of the Arp2/3 complex at actin branch junctions. PLOS Biol 3:e383.

Emerson JJ, Kaessmann H, Betran E, Long M. 2004. Extensive gene traffic on the mammalian X chromosome. Science 303:537–540.

Frankel S, Mooseker MS. 1996. The actin-related proteins. Curr Opin Cell Biol 8:30–37.

Fu J, Wang Y, Fok KL, Yang D, Qiu Y, Chan HC, Koide SS, Miao S, Wang L. 2012. Anti-ACTL7a antibodies: a cause of infertility. Fertil Steril 97:1226–1233 e1221-1228.

Fyrberg C, Ryan L, Kenton M, Fyrberg E. 1994. Genes encoding actin-related proteins of Drosophila melanogaster. J Mol Biol 241:498–503.

Gallach M, Betran E. 2011. Intralocus sexual conflict resolved through gene duplication. Trends Ecol Evol 26:222–228.

Gandhi M, Smith BA, Bovellan M, Paavilainen V, Daugherty-Clarke K, Gelles J, Lappalainen P, Goode BL. 2010. GMF is a cofilin homolog that binds Arp2/3 complex to stimulate filament debranching and inhibit actin nucleation. Curr Biol 20:861–867.

Goodson HV, Hawse WF. 2002. Molecular evolution of the actin family. J Cell Sci 115:2619–2622.

Gramates LS, Marygold SJ, Santos GD, Urbano JM, Antonazzo G, Matthews BB, Rey AJ, Tabone CJ, Crosby MA, Emmert DB, et al. 2017. FlyBase at 25: looking to the future. Nucleic Acids Res 45:D663–D671.

Guindon S, Dufayard JF, Lefort V, Anisimova M, Hordijk W, Gascuel O. 2010. New algorithms and methods to estimate maximum-likelihood phylogenies: assessing the performance of PhyML 3.0. Syst Biol 59:307–321.

Hammesfahr B, Kollmar M. 2012. Evolution of the eukaryotic dynactin complex, the activator of cytoplasmic dynein. BMC Evol Biol 12:95.

Hara Y, Yamagata K, Oguchi K, Baba T. 2008. Nuclear localization of profilin III-ArpM1 complex in mouse spermiogenesis. FEBS Lett 582:2998–3004.

Harata M, Oma Y, Tabuchi T, Zhang Y, Stillman DJ, Mizuno S. 2000. Multiple actin-related proteins of Saccharomyces cerevisiae are present in the nucleus. J Biochem 128:665–671.

Heid H, Figge U, Winter S, Kuhn C, Zimbelmann R, Franke W. 2002. Novel actin-related proteins Arp-T1 and Arp-T2 as components of the cytoskeletal calyx of the mammalian sperm head. Exp Cell Res 279:177–187.

Hocky GM, Baker JL, Bradley MJ, Sinitskiy AV, De La Cruz EM, Voth GA. 2016. Cations Stiffen Actin Filaments by Adhering a Key Structural Element to Adjacent Subunits. J Phys Chem B 120:4558–4567.

Holland DO, Johnson ME. 2018. Stoichiometric balance of protein copy numbers is measurable and functionally significant in a protein-protein interaction network for yeast endocytosis. PLoS Comput Biol 14:e1006022.

Holman L, Freckleton RP, Snook RR. 2008. What use is an infertile sperm? A comparative study of sperm-heteromorphic Drosophila. Evolution 62:374–385.

Holman L, Snook RR. 2006. Spermicide, cryptic female choice and the evolution of sperm form and function. J Evol Biol 19:1660–1670.

Holman L, Snook RR. 2008. A sterile sperm caste protects brother fertile sperm from female-mediated death in Drosophila pseudoobscura. Curr Biol 18:292–296.

Izore T, Kureisaite-Ciziene D, McLaughlin SH, Lowe J. 2016. Crenactin forms actin-like double helical filaments regulated by arcadin-2. Elife 5.

Jagadeeshan S, Singh RS. 2005. Rapidly evolving genes of Drosophila: differing levels of selective pressure in testis, ovary, and head tissues between sibling species. Mol Biol Evol 22:1793–1801.

Jiang X, Assis R. 2017. Natural Selection Drives Rapid Functional Evolution of Young Drosophila Duplicate Genes. Mol Biol Evol 34:3089–3098.

Joly D, Cariou, M. L., Lachaise, D. & David, J. R. 1989. Variation of sperm length and heteromorphism in Drosophilid species. Genetics Selection and Evolution 21:283–293.

Joly D, Cariou, M.L. & Lachaise, D. 1991. Can sperm competition explain sperm polymorphism in Drosophila teissieri? EvolucioUn BioloUgica 5:25–44.

Joly DL D. 1994. Polymorphism in the sperm heteromorphic species of the Drosophila obscura group. Journal of Insect Physiology 40:933–938.

Kabsch W, Mannherz HG, Suck D, Pai EF, Holmes KC. 1990. Atomic structure of the actin:DNase I complex. Nature 347:37–44.

Kaessmann H. 2010. Origins, evolution, and phenotypic impact of new genes. Genome Res 20:1313–1326.

Katkar HH, Davtyan A, Durumeric AEP, Hocky GM, Schramm AC, De La Cruz EM, Voth GA. 2018. Insights into the Cooperative Nature of ATP Hydrolysis in Actin Filaments. Biophys J 115:1589–1602.

Katoh K, Standley DM. 2013. MAFFT multiple sequence alignment software version 7: improvements in performance and usability. Mol Biol Evol 30:772–780.

Kearse M, Moir R, Wilson A, Stones-Havas S, Cheung M, Sturrock S, Buxton S, Cooper A, Markowitz S, Duran C, et al. 2012. Geneious Basic: an integrated and extendable desktop software platform for the organization and analysis of sequence data. Bioinformatics 28:1647–1649.

Kelley LA, Mezulis S, Yates CM, Wass MN, Sternberg MJ. 2015. The Phyre2 web portal for protein modeling, prediction and analysis. Nat Protoc 10:845–858.

Klages-Mundt NL, Kumar A, Zhang Y, Kapoor P, Shen X. 2018. The Nature of Actin-Family Proteins in Chromatin-Modifying Complexes. Front Genet 9:398.

Kleene KC. 2005. Sexual selection, genetic conflict, selfish genes, and the atypical patterns of gene expression in spermatogenic cells. Dev Biol 277:16–26.

Korbie DJ, Mattick JS. 2008. Touchdown PCR for increased specificity and sensitivity in PCR amplification. Nat Protoc 3:1452–1456.

Kosakovsky Pond SL, Posada D, Gravenor MB, Woelk CH, Frost SD. 2006. GARD: a genetic algorithm for recombination detection. Bioinformatics 22:3096–3098.

Lee IH, Kumar S, Plamann M. 2001. Null mutants of the neurospora actin-related protein 1 pointed-end complex show distinct phenotypes. Mol Biol Cell 12:2195–2206.

Machesky LM, Atkinson SJ, Ampe C, Vandekerckhove J, Pollard TD. 1994. Purification of a cortical complex containing two unconventional actins from Acanthamoeba by affinity chromatography on profilin-agarose. J Cell Biol 127:107–115.

McDonald JH, Kreitman M. 1991. Adaptive protein evolution at the Adh locus in Drosophila. Nature 351:652–654.

Muhua L, Karpova TS, Cooper JA. 1994. A yeast actin-related protein homologous to that in vertebrate dynactin complex is important for spindle orientation and nuclear migration. Cell 78:669–679.

Muller J, Oma Y, Vallar L, Friederich E, Poch O, Winsor B. 2005. Sequence and comparative genomic analysis of actin-related proteins. Mol Biol Cell 16:5736–5748.

Mullins RD, Heuser JA, Pollard TD. 1998. The interaction of Arp2/3 complex with actin: nucleation, high affinity pointed end capping, and formation of branching networks of filaments. Proc Natl Acad Sci U S A 95:6181–6186.

Noguchi T, Lenartowska M, Rogat AD, Frank DJ, Miller KG. 2008. Proper cellular reorganization during Drosophila spermatid individualization depends on actin structures composed of two domains, bundles and meshwork, that are differentially regulated and have different functions. Mol Biol Cell 19:2363–2372.

Noguchi T, Miller KG. 2003. A role for actin dynamics in individualization during spermatogenesis in Drosophila melanogaster. Development 130:1805–1816.

Nolen BJ, Tomasevic N, Russell A, Pierce DW, Jia Z, McCormick CD, Hartman J, Sakowicz R, Pollard TD. 2009. Characterization of two classes of small molecule inhibitors of Arp2/3 complex. Nature 460:1031–1034.

Oda T, Iwasa M, Aihara T, Maeda Y, Narita A. 2009. The nature of the globular- to fibrous-actin transition. Nature 457:441–445.

Oppliger A, Hosken, D.J. & Ribi, G. 1998. Snail sperm production characteristics vary with sperm competition risk. Proc. R. Soc. Lond. B 265:1527–1534.

Peterson CL, Zhao Y, Chait BT. 1998. Subunits of the yeast SWI/SNF complex are members of the actin-related protein (ARP) family. J Biol Chem 273:23641–23644.

Pfaendtner J, Volkmann N, Hanein D, Dalhaimer P, Pollard TD, Voth GA. 2012. Key structural features of the actin filament Arp2/3 complex branch junction revealed by molecular simulation. J Mol Biol 416:148–161.

Phillips JC, Braun R, Wang W, Gumbart J, Tajkhorshid E, Villa E, Chipot C, Skeel RD, Kale L, Schulten K. 2005. Scalable molecular dynamics with NAMD. J Comput Chem 26:1781–1802.

Pollard TD. 2007. Regulation of actin filament assembly by Arp2/3 complex and formins. Annu Rev Biophys Biomol Struct 36:451–477.

Pollard TD, Borisy GG. 2003. Cellular motility driven by assembly and disassembly of actin filaments. Cell 112:453–465.

Rouiller I, Xu XP, Amann KJ, Egile C, Nickell S, Nicastro D, Li R, Pollard TD, Volkmann N, Hanein D. 2008. The structural basis of actin filament branching by the Arp2/3 complex. J Cell Biol 180:887–895.

Russo CAMM, Beatriz; Frazão, Annelise; Voloch, Carolina M. 2013. Phylogenetic analysis and a time tree for a large drosophilid data set (Diptera: Drosophilidae). Zoological Journal of the Linnean Society 169:765–775.

Saunders MG, Voth GA. 2012. Comparison between actin filament models: coarse-graining reveals essential differences. Structure 20:641–653.

Schafer DA, Gill SR, Cooper JA, Heuser JE, Schroer TA. 1994. Ultrastructural analysis of the dynactin complex: an actin-related protein is a component of a filament that resembles F-actin. J Cell Biol 126:403–412.

Smith NG, Eyre-Walker A. 2002. Adaptive protein evolution in Drosophila. Nature 415:1022–1024.

Snook RR, Markow TA, Karr TL. 1994. Functional nonequivalence of sperm in Drosophila pseudoobscura. Proc Natl Acad Sci U S A 91:11222–11226.

Suyama M, Torrents D, Bork P. 2006. PAL2NAL: robust conversion of protein sequence alignments into the corresponding codon alignments. Nucleic Acids Res 34:W609–612.

Swallow JG, Wilkinson GS. 2002. The long and short of sperm polymorphisms in insects. Biol Rev Camb Philos Soc 77:153–182.

Tanaka H, Iguchi N, Egydio de Carvalho C, Tadokoro Y, Yomogida K, Nishimune Y. 2003. Novel actin-like proteins T-ACTIN 1 and T-ACTIN 2 are differentially expressed in the cytoplasm and nucleus of mouse haploid germ cells. Biol Reprod 69:475–482.

Thibault ST, Singer MA, Miyazaki WY, Milash B, Dompe NA, Singh CM, Buchholz R, Demsky M, Fawcett R, Francis-Lang HL, et al. 2004. A complementary transposon tool kit for Drosophila melanogaster using P and piggyBac. Nat Genet 36:283–287.

Turner LM, Chuong EB, Hoekstra HE. 2008. Comparative analysis of testis protein evolution in rodents. Genetics 179:2075–2089.

van den Ent F, Amos LA, Lowe J. 2001. Prokaryotic origin of the actin cytoskeleton. Nature 413:39–44.

Vibranovski MD, Lopes HF, Karr TL, Long M. 2009. Stage-specific expression profiling of Drosophila spermatogenesis suggests that meiotic sex chromosome inactivation drives genomic relocation of testis-expressed genes. PLoS Genet 5:e1000731.

Vibranovski MD, Zhang Y, Long M. 2009. General gene movement off the X chromosome in the Drosophila genus. Genome Res 19:897–903.

Vinckenbosch N, Dupanloup I, Kaessmann H. 2006. Evolutionary fate of retroposed gene copies in the human genome. Proc Natl Acad Sci U S A 103:3220–3225.

Waterhouse A, Bertoni M, Bienert S, Studer G, Tauriello G, Gumienny R, Heer FT, de Beer TAP, Rempfer C, Bordoli L, et al. 2018. SWISS-MODEL: homology modelling of protein structures and complexes. Nucleic Acids Res 46:W296–W303.

Yang Z. 2007. PAML 4: phylogenetic analysis by maximum likelihood. Mol Biol Evol 24:1586–1591.

Zhou Q, Bachtrog D. 2012. Sex-specific adaptation drives early sex chromosome evolution in Drosophila. Science 337:341–345.

