## Supplementary Figures 1-12 for "A burst of genetic innovation in actin-related proteins (Arps) for testis-specific function in a *Drosophila* lineage"

### Supplemental Figure S1: Sequenced *Drosophila* species encode canonical Arps and lineage-specific Arps.

**A**

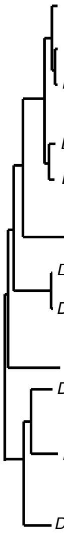

| | Actin | Dup 1 | Dup 2 | Dup 3 | Arp1 | Arp2 | Arp2D | Arp3 | Arp4 | Arp5 | Arp6 | Arp8 | Arp10 |
|---|---|---|---|---|---|---|---|---|---|---|---|---|---|
| *D. melanogaster* | 2 | 0 | 0 | 0 | 1 | 1 | 0 | 1 | 1 | 1 | 1 | 1 | 1 |
| *D. simulans* | 2 | 0 | 0 | 0 | 1 | 1 | 0 | 1 | 1 | 1 | 1 | 1 | 1 |
| *D. sechellia* | 2 | 0 | 0 | 0 | 1 | 1 | 0 | 1 | 1 | 1 | 1 | 1 | 1 |
| *D. yakuba* | 2 | 0 | 0 | 0 | 1 | 1 | 0 | 1 | 1 | 1 | 1 | 1 | 1 |
| *D. erecta* | 2 | 0 | 0 | 0 | 1 | 1 | 0 | 1 | 1 | 1 | 1 | 1 | 1 |
| *D. ananassae* | 2 | 0 | 0 | 0 | 1 | 1 | 0 | 1 | 1 | 1 | 1 | 1 | 1 |
| *D. pseudoobscura* | 2 | 1 | 1 | 1 | 1 | 1 | 1 | 1 | 1 | 1 | 1 | 1 | 1 |
| *D. persimilis* | 2 | 1 | 1 | 1 | 1 | 1 | 1 | 1 | 1 | 1 | 1 | 1 | 1 |
| *D. willistoni* | 2 | 0 | 0 | 0 | 1 | 1 | 0 | 1 | 1 | 1 | 1 | 1 | 1 |
| *D. virilis* | 2 | 0 | 0 | 0 | 1 | 1 | 0 | 1 | 1 | 1 | 1 | 1 | 1 |
| *D. mojavensis* | 2 | 0 | 0 | 0 | 1 | 1 | 0 | 1 | 1 | 1 | 1 | 1 | 1 |
| *D. grimshawi* | 2 | 0 | 0 | 0 | 1 | 1 | 0 | 1 | 1 | 1 | 1 | 1 | 1 |

**B**

| | Gene | Chromosomal location | |
|---|---|---|---|
| | | ***D. mel*** | ***D. pse*** |
| Actin | **Act5C** | X | XL_group1e |
| | **Act42A** | 2R | 3 |
| Muscle Actin | **Act57B** | 2R | 3 |
| | **Act87E** | 3R | 2 |
| | **Act88F** | 3R | 2 |
| | **Act79B** | 3L | XR_grp8 |
| | **Arp2** | X | XL_grp1e |
| Duplicates | **Dup 1** | NA | 4_grp4 |
| | **Dup 2** | NA | 4_grp3 |
| | **Dup 3** | NA | XL_grp1a |
| | **Arp2D** | NA | 4_grp3 |

**C**

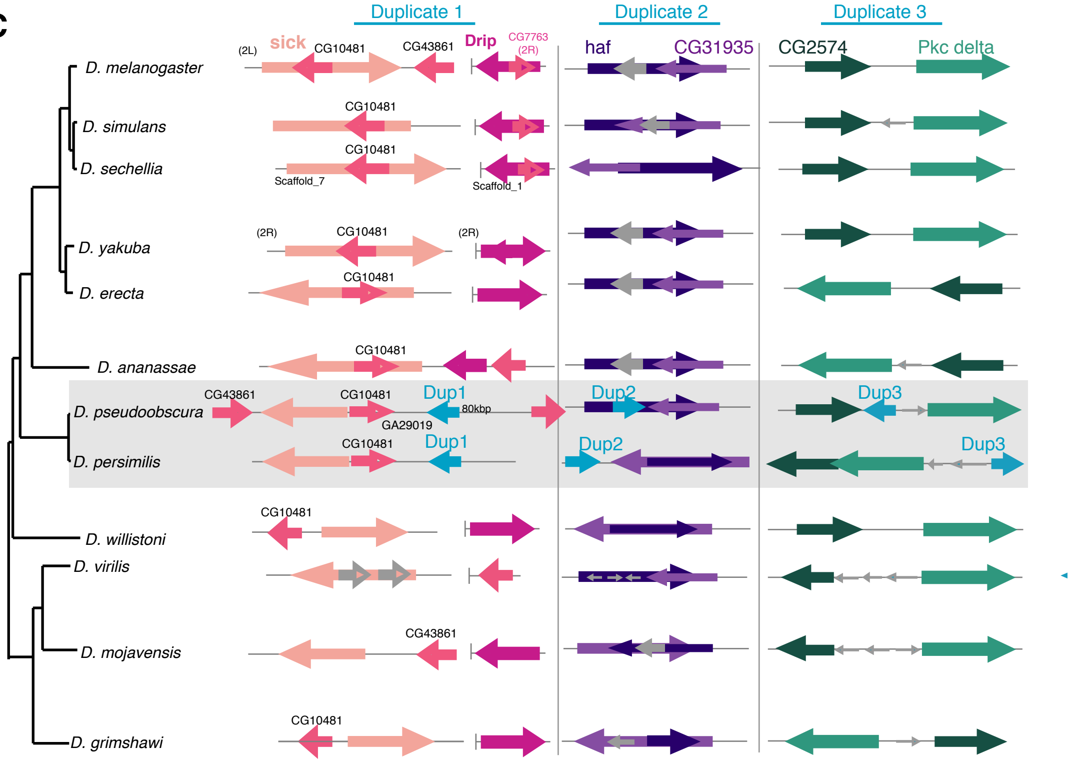

**Supplemental Figure S2: Arp2D is present in only the obscura clade.**

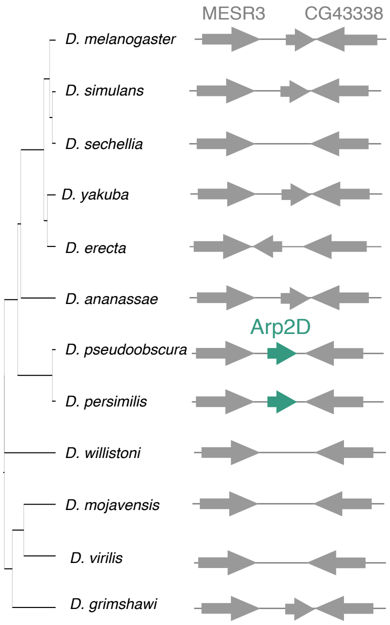

**Supplemental Figure S3: Nucleotide trees including pseudogenized genes recapitulate overall species tree topology**

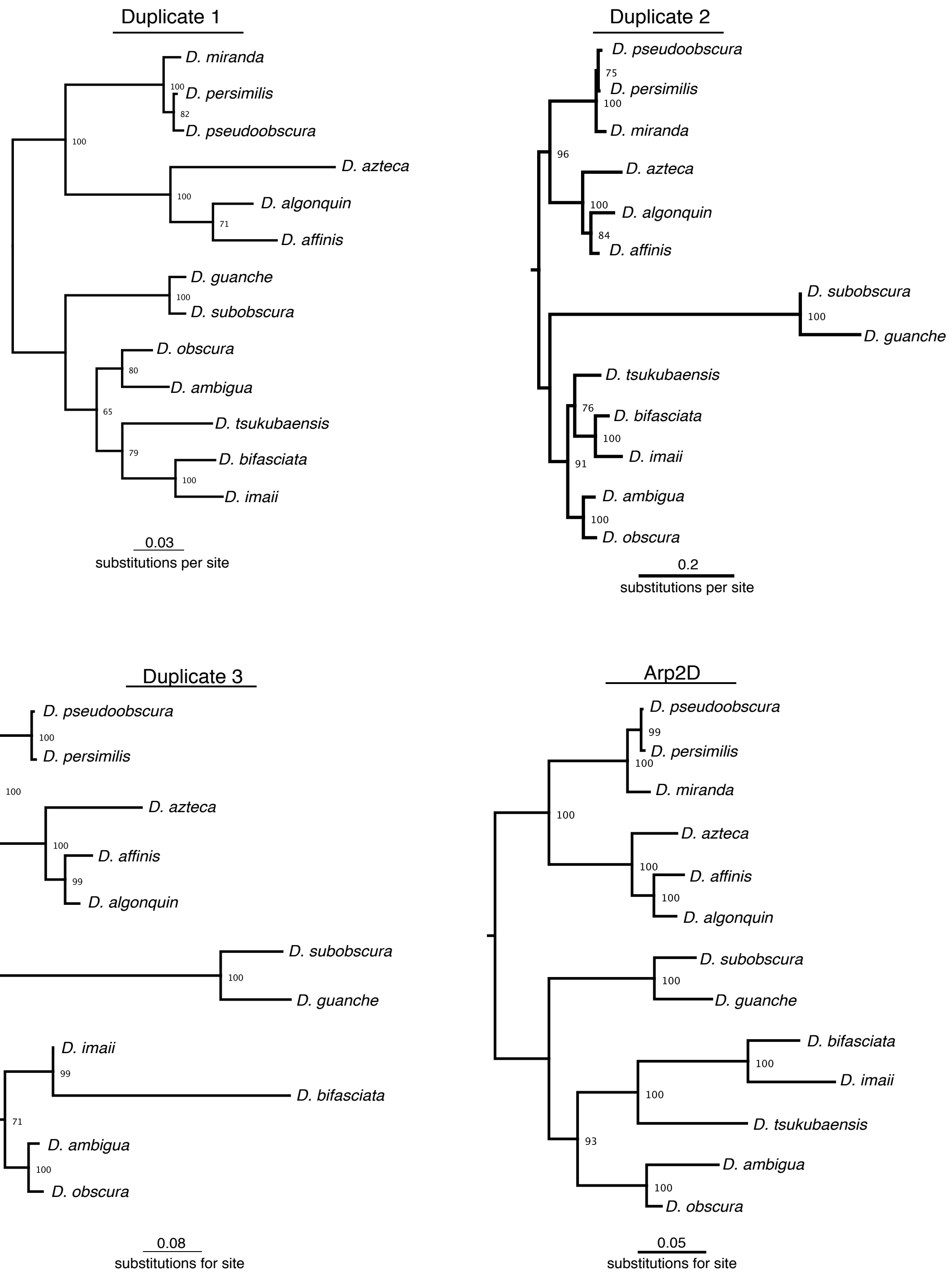

### Supplemental Figure S4: A single base-pair deletion leads to pseudogenization of Arp2D in *D. subobscura*

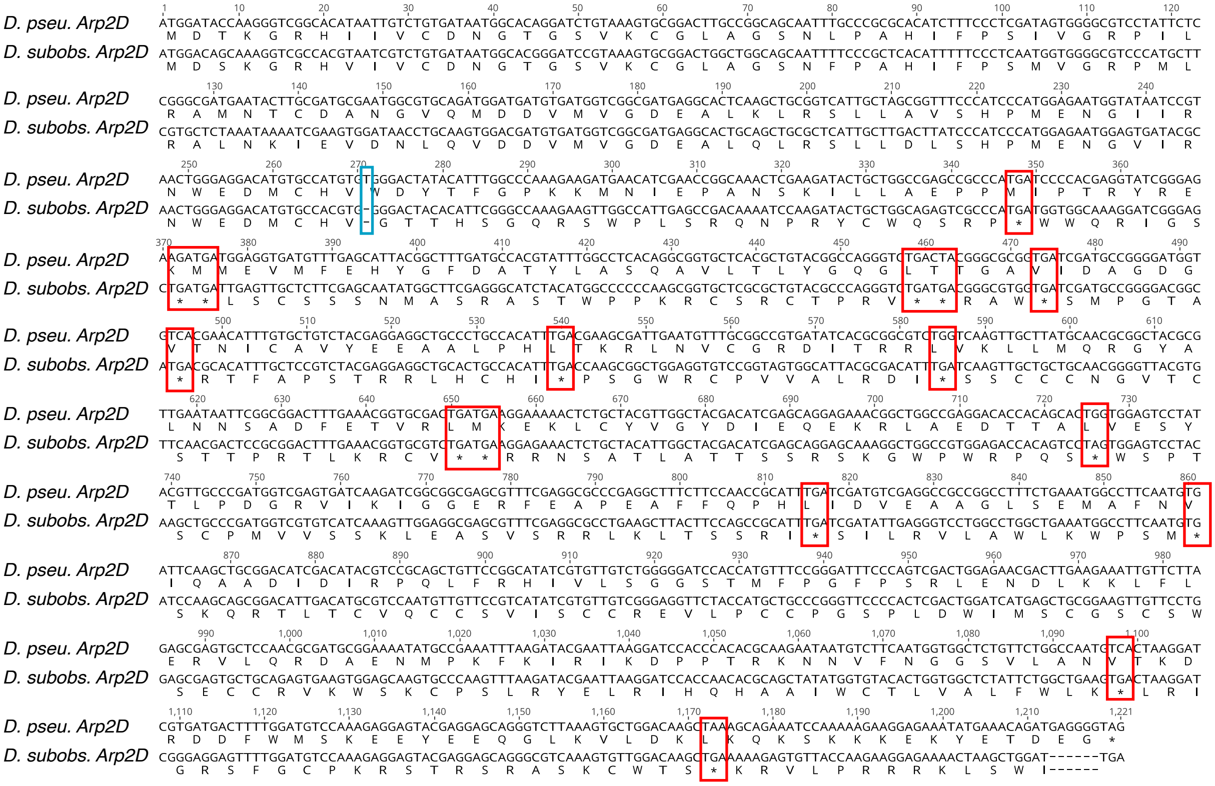

### Supplemental Figure S5: *D. miranda* encodes full-length Dup2 and Arp2D.

**A**

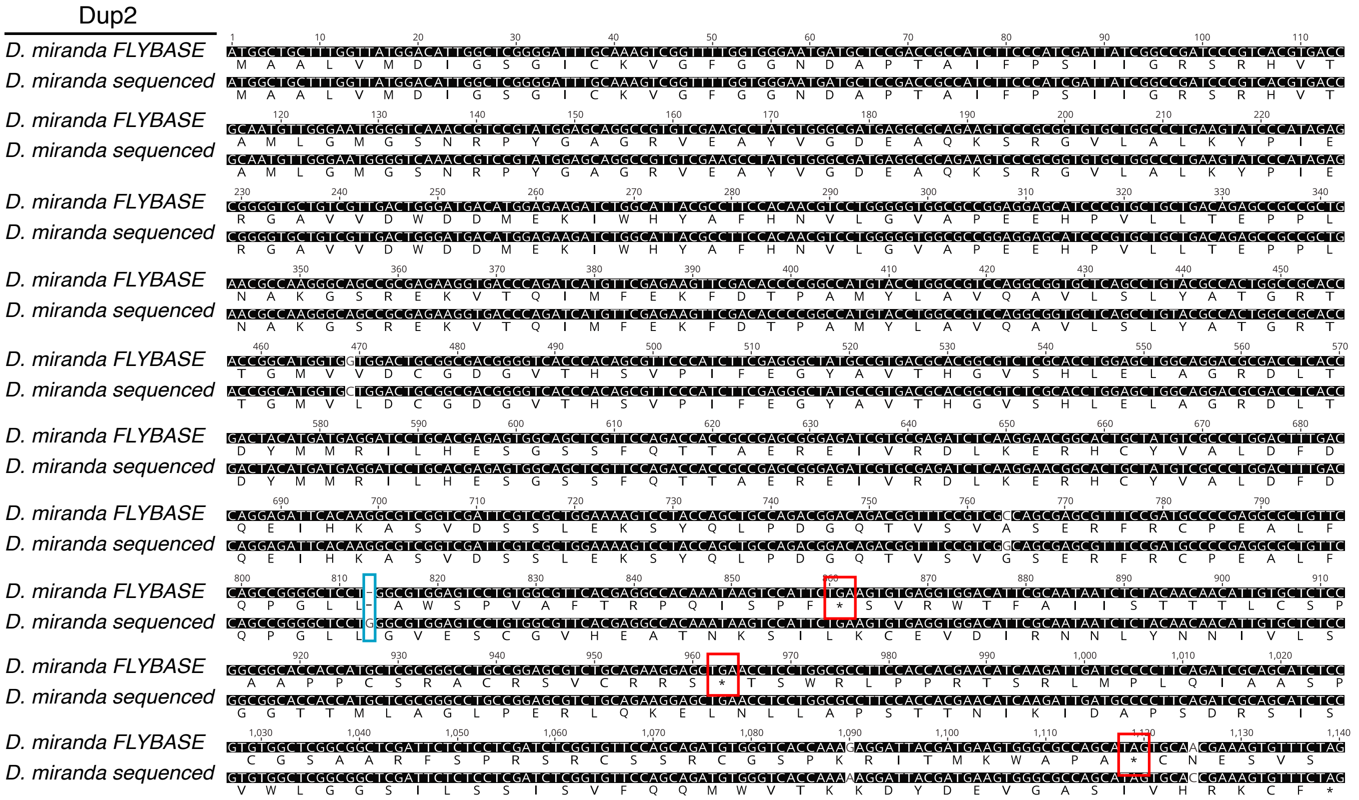


**B**

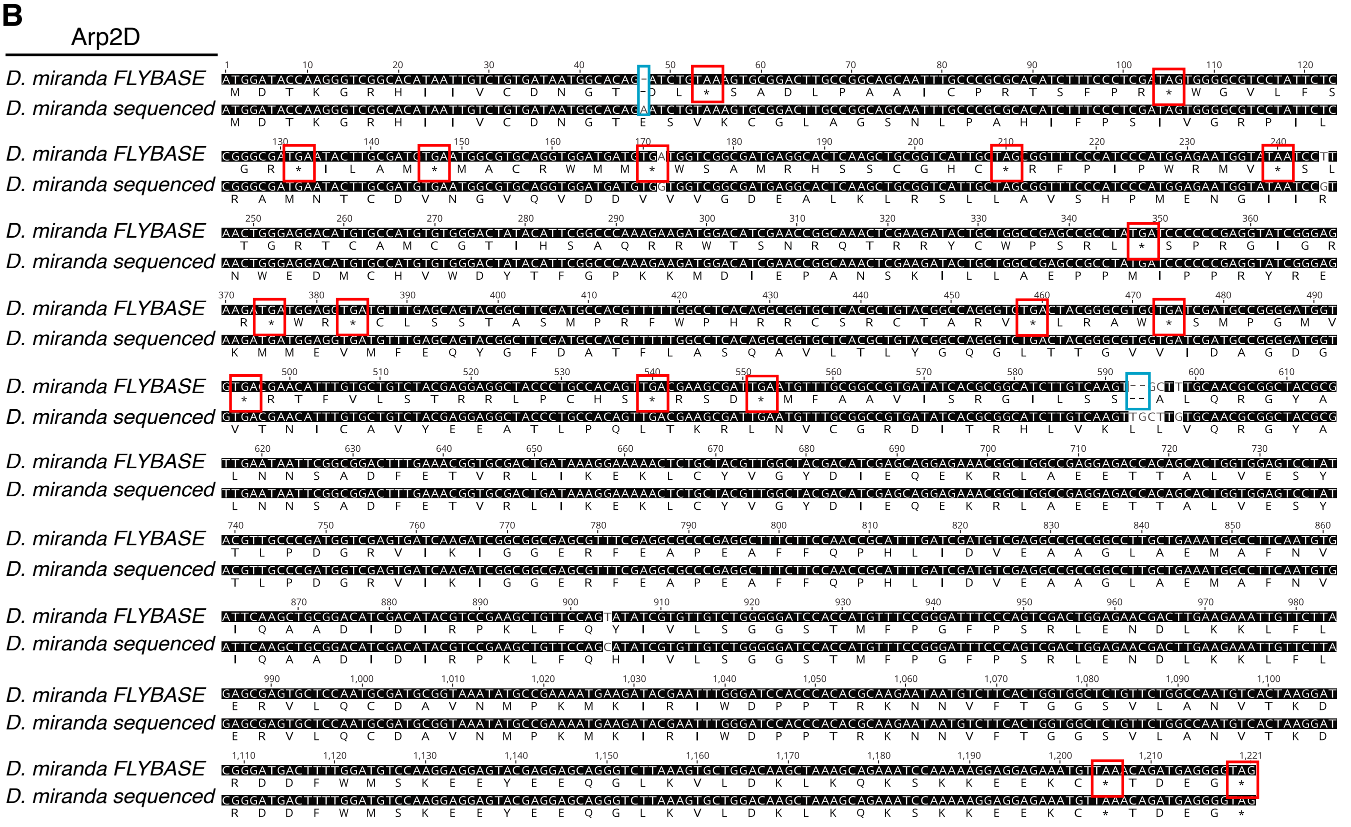

**Supplementary Figure S6: Gene and protein trees of Duplicates 1-3 and canonical Arps**

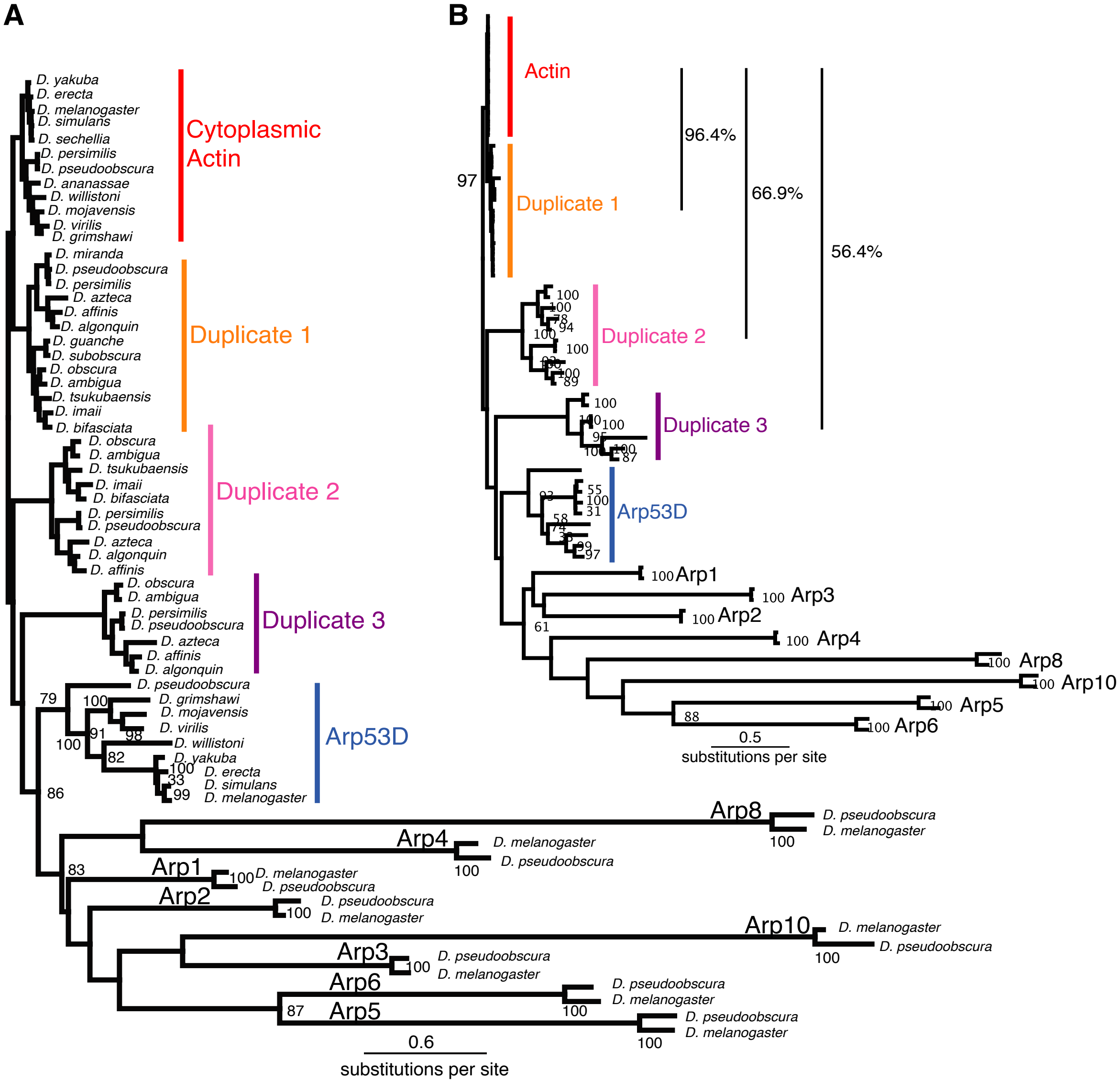

### Supplemental Figure S7: RT-PCR indicates male-enriched expression of Arp duplicates

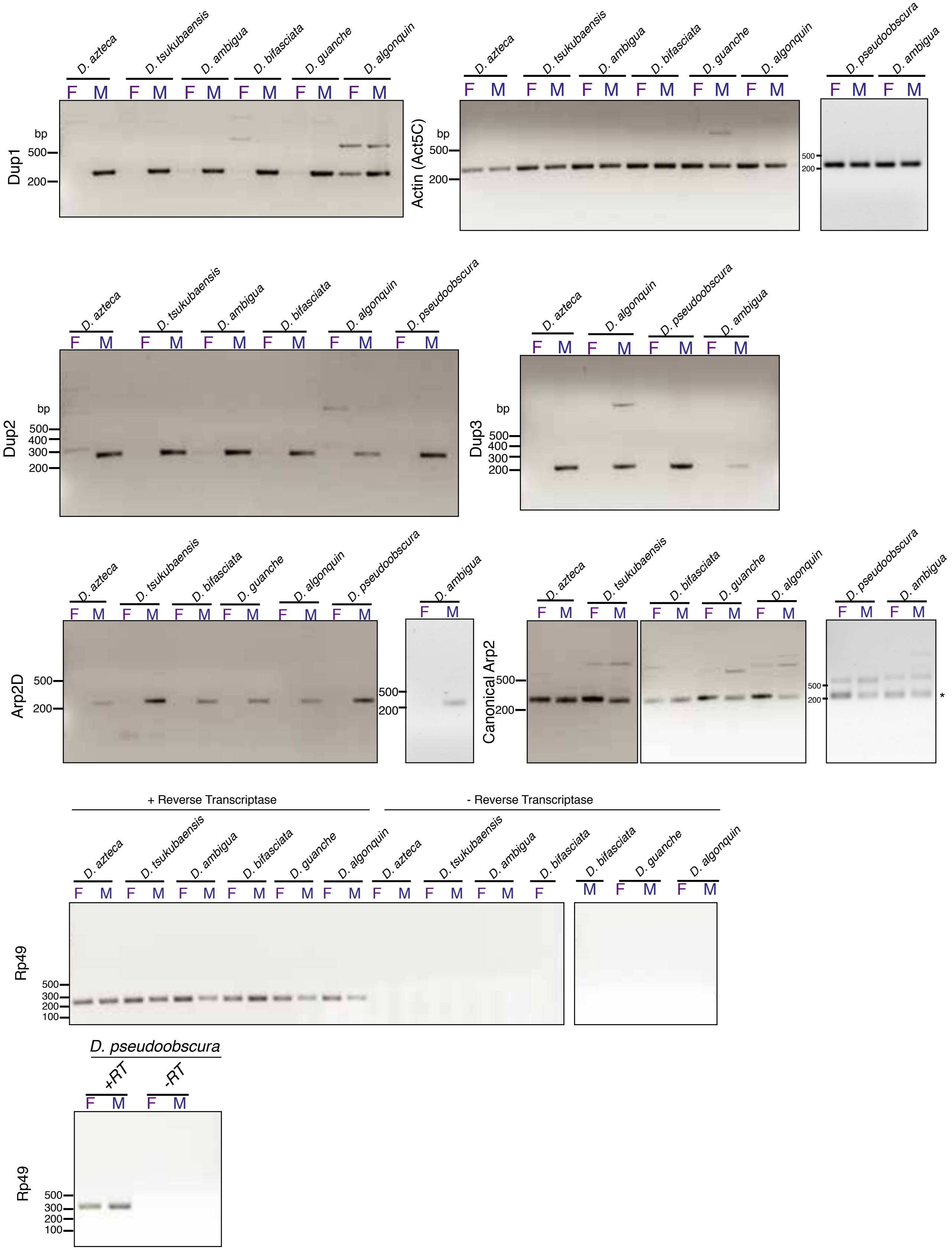

**Supplemental Figure S8: The ATP-binding motifs are conserved in the *D. pseudoobscura* Arp duplicates**

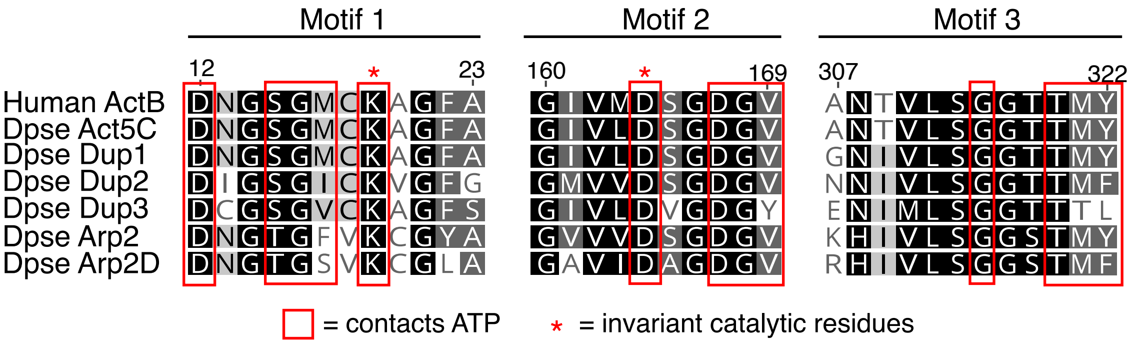

**Supplemental Figure S9: Arp2D/3 exhibits a stable complex similar to Arp2/3 despite fixed residue changes**

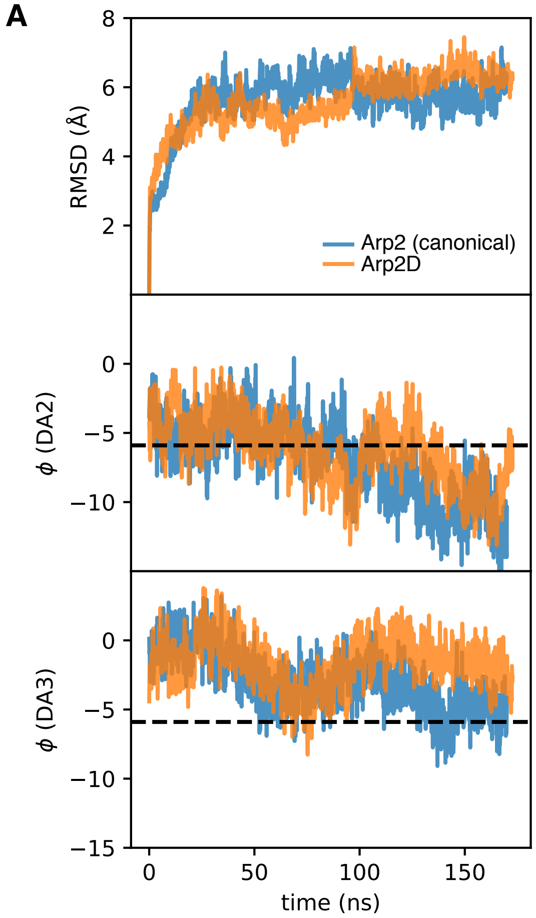


B

| Arp2 residue | Arp2D residue |
|---|---|
| E51 | Q53 |
| S67 | L64 |
| Y22 | L22 |
| A364 | S361 |
| R354 | T351 |
| S164 | A161 |
| T290 | V287 |
| L270 | F268 |
| M234 | Q231 |

**Supplemental Figure S10: Full-length Arp2D-sfGFP protein is expressed in the testis.**

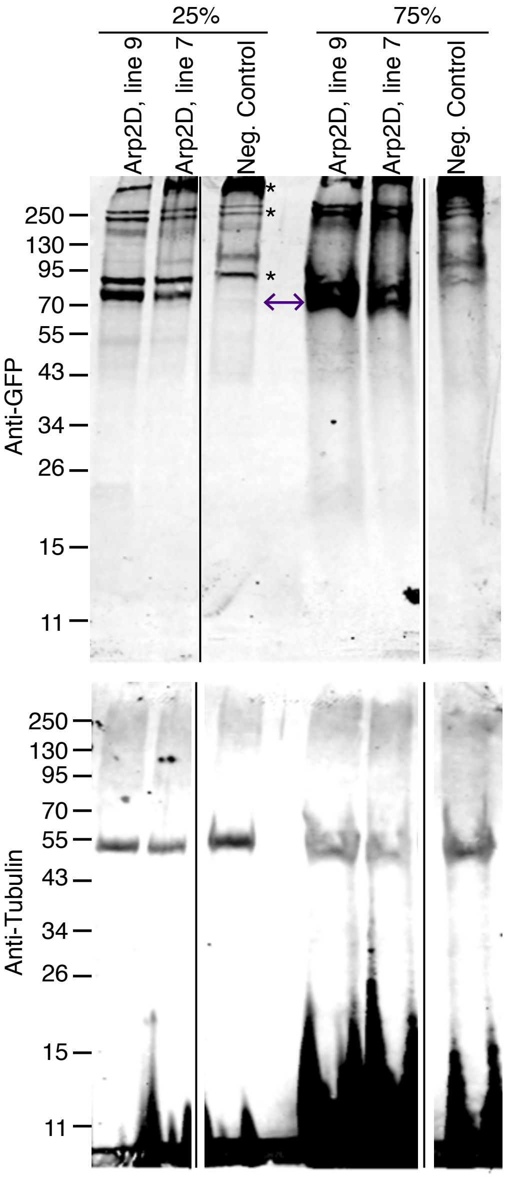

**Supplemental Figure S11: *D. pseudoobscura* w- mature sperm exhibit autofluorescence but none in meiotic and post-meiotic cysts**

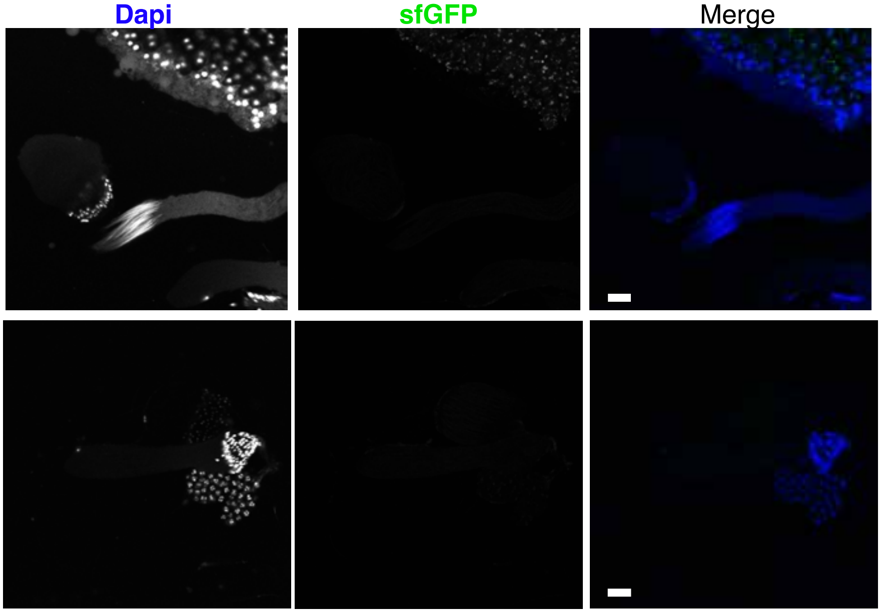

### Supplemental Figure S12: Arp2D localizes only to motile fan-like actin cones

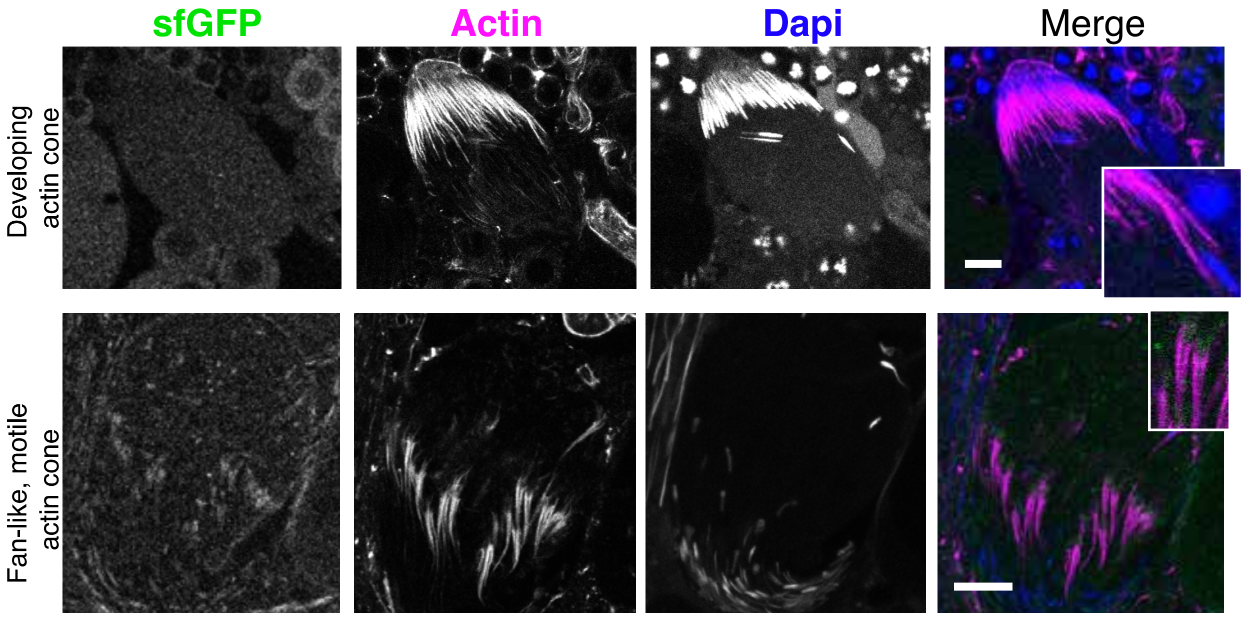
